# Invertebrate methylomes provide insight into mechanisms of environmental tolerance and reveal methodological biases

**DOI:** 10.1101/2021.03.29.437539

**Authors:** Shelly A. Trigg, Yaamini R. Venkataraman, Mackenzie R. Gavery, Steven B. Roberts, Debashish Bhattacharya, Alan Downey-Wall, Jose M. Eirin-Lopez, Kevin M. Johnson, Katie E. Lotterhos, Jonathan B. Puritz, Hollie M. Putnam

**Affiliations:** University of Washington, School of Aquatic and Fishery Sciences 1122 NE Boat St. Seattle, WA, 98195, USA; Environmental and Fisheries Sciences Division, Northwest Fisheries Science Center, National Marine Fisheries Service, National Oceanic and Atmospheric Administration, 2725 Montlake Blvd E, Seattle, WA, 98112, USA; Department of Biochemistry and Microbiology, Rutgers University, New Brunswick, NJ 08901 USA; Department of Marine and Environmental Sciences, Northeastern University, 430 Nahant Road, Nahant, MA 01908; Florida International University, Environmental Epigenetics Laboratory, Institute of Environment 3000 NE 151 St. North Miami, FL, 33181, USA; Center for Coastal Marine Sciences, California Polytechnic State University, San Luis Obispo, CA, 93407, USA; California Sea Grant, University of California San Diego, La Jolla, CA, 92093; Department of Biological Sciences, University of Rhode Island, Kingston, RI 02881, USA

**Keywords:** bisulfite sequencing, coral, epigenetics, marine invertebrate

## Abstract

There is a growing focus on the role of DNA methylation in the ability of marine invertebrates to rapidly respond to changing environmental factors and anthropogenic impacts. However, genome-wide DNA methylation studies in non-model organisms are currently hampered by limited understanding of methodological biases. Here we compare three methods for quantifying DNA methylation at single base-pair resolution — Whole Genome Bisulfite Sequencing (WGBS), Reduced Representation Bisulfite Sequencing (RRBS), and Methyl-CpG Binding Domain Bisulfite Sequencing (MBDBS) — using multiple individuals from two reef-building coral species with contrasting environmental sensitivity. All methods reveal substantially greater methylation in *Montipora capitata* (11.4%) than the more sensitive *Pocillopora acuta* (2.9%). The majority of CpG methylation in both species occurs in gene bodies and flanking regions. In both species, MBDBS has the greatest capacity for detecting CpGs in coding regions at our sequencing depth, however MBDBS may be influenced by intra-sample methylation heterogeneity. RRBS yields robust information for specific loci albeit without enrichment of any particular genome feature and with significantly reduced genome coverage. Relative genome size strongly influences the number and location of CpGs detected by each method when sequencing depth is limited, illuminating nuances in cross-species comparisons. As genome-wide methylation differences, supported by data across bisulfite sequencing methods, may contribute to environmental sensitivity phenotypes in critical marine invertebrate taxa, these data provide a genomic resource for investigating the functional role of DNA methylation in environmental tolerance.

## Introduction

Environmental stimuli interact with genomic content to drive variation in gene and protein expression, resulting in phenotypic plasticity. This plasticity has the potential to buffer against mortality under environmental change (Baldwin 1902), or conversely be maladaptive (Velotta et al. 2018). Furthermore, plasticity may enhance or diminish evolutionary rates (Ghalambor et al. 2007), which is particularly relevant to plasticity-evolution feedbacks (Ghalambor et al. 2015, 2007; Kronholm and Collins 2016). This is of particular concern in the Anthropocene (Lewis and Maslin 2015), as global change exacerbates the mismatch between phenotype and a rapidly changing environment.

The increase in negative global climate change consequences have prompted an intensification of research into phenotypic plasticity, gene regulation, and epigenetic mechanisms in non-model marine invertebrates [as reviewed in (Eirin-Lopez and Putnam 2019; Roberts and Gavery 2012; Hofmann 2017)]. Specifically, carryover effects and cross- and multi-generational plasticity in response to climate change (Byrne et al. 2020) may be generated by epigenetic regulation of gene expression (Liew et al. 2020, 2018; Dixon et al. 2018). As epigenetic research has increased there has been a focus on DNA methylation: the addition of a methyl group on the cytosine residues in the genome often in the cytosine phosphate guanine (CpG) context (Zemach et al. 2010). DNA methylation has gene expression regulation capacity through the interaction of base modification with transcriptional elements. Early bulk enzyme-based and fingerprinting methods for quantifying DNA methylation in marine invertebrates provided initial insights into DNA methylation and organismal phenotypic plasticity in response to environmental changes (Putnam, Davidson, and Gates 2016; Gavery and Roberts 2010; Rodriguez-Casariego et al. 2018; Dimond, Gamblewood, and Roberts 2017; Suarez-Ulloa et al. 2018; Gonzalez-Romero et al. 2017; Riviere et al. 2013).

Non-sequencing approaches that quantify global or bulk DNA methylation [e.g., colorimetric or fluorescent ELISAs (Putnam, Davidson, and Gates 2016; Gavery and Roberts 2010; Rodriguez-Casariego et al. 2018; Dimond, Gamblewood, and Roberts 2017; Riviere et al. 2013) are low-cost, rapidly applied, and do not require genomic resources to generate information on the responsiveness of the methylome. These global estimates do not, however, fully capture local changes in DNA methylation across different genome regions. Specifically, differences in the location and amount of methylation in two samples or treatments could lead to biased conclusions when based on average percent methylation at the bulk level. Consequently, non-sequencing methods are limited in their ability to elucidate specific mechanisms of expression regulation and thus are unable to fully address the functional implications of methylation-driven regulation within the genome. In contrast, the use of genome-wide approaches that provide single base-pair resolution allow the testing of hypotheses regarding spurious transcription, alternative splicing, and exon skipping (Roberts and Gavery 2012). For example, the use of Whole Genome Bisulfite Sequencing (WGBS) to investigate the role of DNA methylation in regulating genes involved in caste specification in honeybees identified differential methylation in an exon of the *anaplastic lymphoma kinase (ALK)* gene; this exon was differentially retained in a splice variant between queens and workers (Foret et al. 2012). Thus there is a clear need for single base-pair assessment of DNA methylomes facilitated by next generation sequencing to more fully elucidate the relationship of DNA methylation and gene expression in non-model invertebrates.

Genome-wide levels of DNA methylation can be estimated by several bisulfite conversion and sequencing approaches. Bisulfite conversion of DNA results in the deamination of unmethylated cytosine to uracil, which leaves a base change signature in the DNA that can be tracked via sequence comparison between bisulfite-converted samples and reference genomes. While the number of bisulfite sequencing approaches are expanding [e.g., epiGBS (van Gurp et al. 2016)], the widely-used approaches are WGBS, Reduced Representation Bisulfite Sequencing (RRBS), and more recently, Methyl-CpG Binding Domain Bisulfite Sequencing (MBDBS). WGBS is considered to be the gold-standard of bisulfite sequencing because it provides full coverage of the genome (given deep sequencing coverage) and the capacity to detect the entire methylome at single base-pair resolution.

While providing a comprehensive approach, the high cost of WGBS is juxtaposed against the often very small fraction of methylated DNA in invertebrate genomes (Tweedie et al. 1997). Alternatively, approaches such as RRBS also use bisulfite conversion to quantitatively assess DNA methylation with base-pair resolution. RRBS incorporates a restriction digestion of the genome to enrich for CpG rich regions, and was designed to enrich for promoters and other genomic regions containing CpG islands because they have important regulatory functions in mammals (Meissner et al. 2008); however, applications of RRBS in lower vertebrates, such as fish, report this method is less biased toward CpG islands (Chatterjee et al. 2013). This is a more cost-effective approach as it only sequences a small portion of the genome, but requires restriction enzyme recognition sites near other CpGs to gather high resolution data. Since DNA methylation in invertebrates is primarily limited to coding regions (Roberts and Gavery 2012; Flores et al. 2012; Dixon et al. 2018), it is less clear whether enrichment of CG-rich DNA using RRBS will enrich for informative or regulatory regions of invertebrate genomes.

In contrast to the CpG-rich, region-specific targeting of RRBS, MBDBS uses Methyl Binding Domain proteins to target and enrich methylated CpGs, then employs bisulfite conversion to provide single base-pair resolution of DNA fragments enriched for methylated regions. Many marine invertebrate genomes consist of highly methylated regions that are distributed in a mosaic pattern throughout predominantly unmethylated DNA (Suzuki et al. 2007). When invertebrate methylomes have been characterized, these highly methylated regions overlap with gene bodies and have been shown to play a role in gene expression activity (Roberts and Gavery 2012). Therefore, using an enrichment approach such as MBDBS to isolate gene body methylation can be a cost-effective and gene body focused alternative to WGBS or RRBS (Gavery and Roberts 2013; Venkataraman et al. 2020). The base-pair resolution and ability to quantify loci methylation offered by the combination of MBD enrichment and BS conversion is an advantage compared to MBD-seq alone (Dixon and Matz 2020), as the latter assumes that methylation level is proportional to read depth. In contrast to WGBS or RRBS, the quantification and interpretation of MBDBS data could be complicated by individual variation in methylation levels (e.g., one individual who has high methylation in a particular region would have data showing enrichment of that region, whereas another individual who lacks methylation in that region would have missing data there).

Given the need to assess plasticity mechanisms and the acclimatization potential of a variety of marine taxa, it is critical to compare the potential of different approaches to to detect, quantify, and assess DNA methylation with respect to specific biological hypotheses of interest. To this end, we studied three DNA methylation quantification approaches that provide single base-pair resolution data using bisulfite conversion and sequencing: WGBS, RRBS, and MBDBS. We applied these methods to two reef-building corals, *Montipora capitata* and *Pocillopora acuta*, which have different environmental sensitivity, phenotypic plasticity, inducible DNA methylation (Putnam, Davidson, and Gates 2016), and genome sizes (Shumaker et al. 2019; Vidal-Dupiol et al. 2019). We assessed species-specific differences in genome-wide methylation and contrasted percent methylation of common loci, gene coverage, and orthologous genes across methods. Then, we compared the coverage and genomic location of CpG data generated from the three methods. Compared to WGBS, both MBDBS and RRBS have advantages and potential limitations associated with biology, genome characteristics, and experimental design, highlighting the need to fully consider these aspects when evaluating DNA methylation for particular hypotheses of methylation function in invertebrates. As part of this effort, we characterized DNA methylation differences in two coral species, providing valuable insights into the epigenetic underpinnings of phenotypic plasticity in non-model marine invertebrates.

## Materials and Methods

### Sample collection

The reef-building scleractinian coral species *Montipora capitata* and *Pocillopora acuta* were collected from 1-2m depth on the patch reefs of Kane ohe Bay Hawai i under SAP 2019-60 between 4 - 7 September 2018. Corals were transported to the Hawai i Institute of Marine Biology where they were held in outdoor tanks under ambient conditions for 15 days, then snap-frozen in liquid nitrogen and stored at −80°C until nucleic acid extraction was performed. For each of the two coral species, fragments were collected from three different individuals in ambient conditions.

### Nucleic Acid Extraction

Samples were removed from −80°C and small tissue fragments were clipped directly into a tube containing RNA/DNA shield (1 ml) and glass beads (0.5 mm). The tissue clippings consisted of all coral cell types and their symbionts. Samples were homogenized on a vortexer for 1 minute for the thin tissue imperforate coral *Pocillopora acuta* and 2 minutes for the thick tissue perforate coral *Montipora capitata* at maximum speed to ensure tissue extraction of all cell types. The supernatant was removed and DNA was extracted using the Zymo Quick-DNA/RNA™ Miniprep Plus Kit and subsequently checked for quality using gel electrophoresis on an Agilent 4200 TapeStation and quantified using a Qubit. One DNA preparation was made from each of the three individuals per coral species and was subsequently divided into three aliquots for each of the three bisulfite sequencing methods (WGBS, MBDBS, and RRBS) to yield a total of 18 libraries (**Figure 1**).

**Figure 1.**
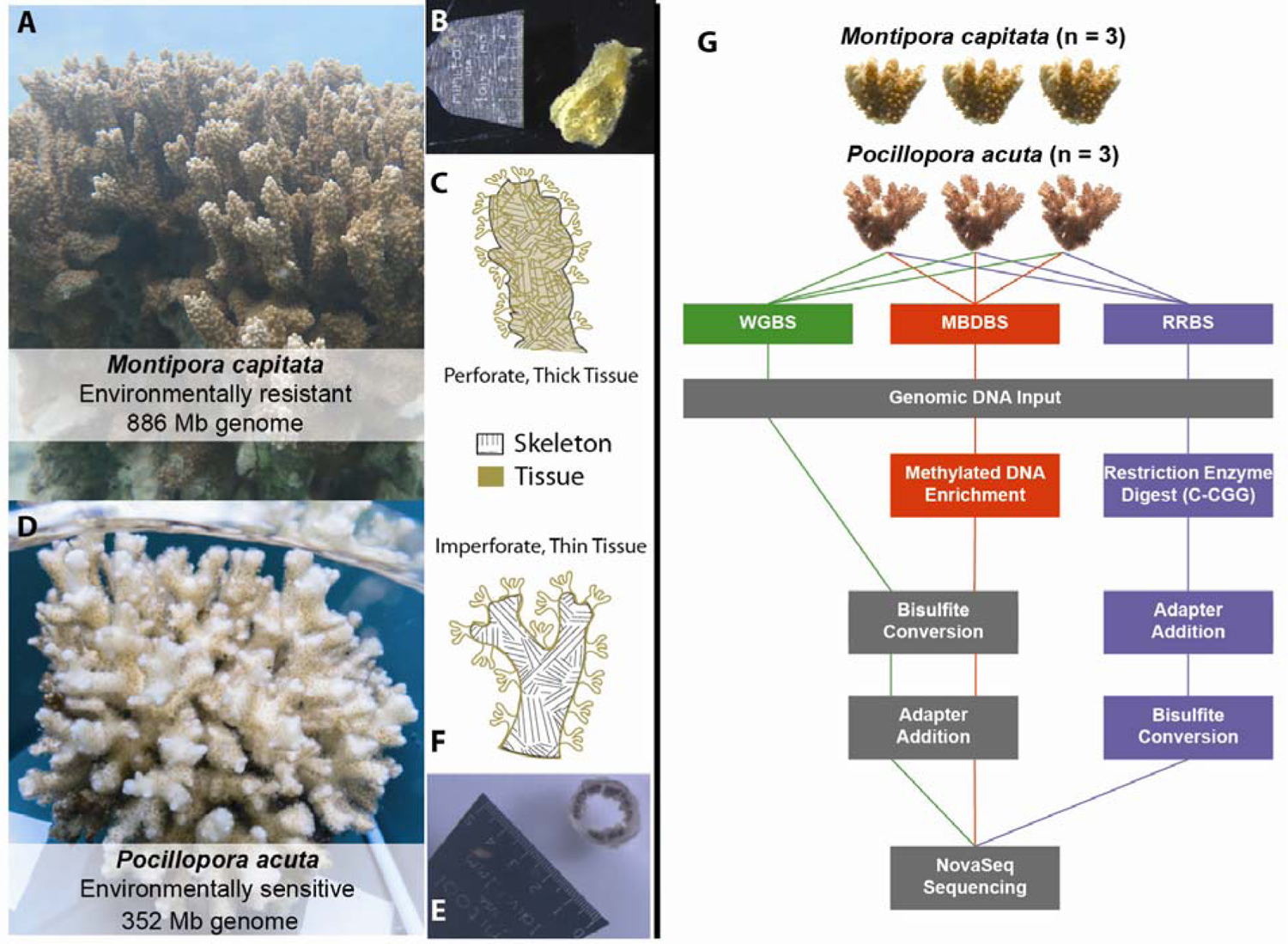
Experimental design. Three biological replicate coral samples were obtained from both coral species. A) *M. capitata*, where B) a cross section of a decalcified fragment reveals thick tissue, and C) a perforate tissue skeletal interaction. In contrast in D) *P. acuta*, a E) a cross section of a decalcified fragment reveals thin tissue, and F) an imperforate tissue skeletal interaction. DNA was extracted from each coral sample and split for use in Whole Genome Bisulfite Sequencing (WGBS), Reduced Representation Bisulfite Sequencing (RRBS), and Methyl-CpG Binding Domain Bisulfite Sequencing (MBDBS) library preparation methods. Three libraries were generated for each of the three methods, yielding nine libraries for each species and 18 libraries total. Tissue photo credit: Ariana Huffmyer. Colony photo credit: Hollie Putnam and Danielle Claar.

### Genome Information

Previously sequenced and assembled coral genomes were used for mapping *Montipora capitata* (Shumaker et al. 2019) and *Pocillopora acuta* (Vidal-Dupiol et al. 2019) DNA methylation data. Both of the coral genomes have a high and similar number of predicted genes (63,227 in *M.capitata* and 64,558 in *P. acuta*). However, *P. acuta* is much smaller in size (∼352MB vs. ∼886 MB in *M. capitata*), has less repetition, a greater number of scaffolds (25,553 in *P. acuta* vs. 3,043 in *M.capitata*), and lower genome assembly continuity (N50 is 171,375 in *P. acuta* and 540,623 in *M. capitata*).

Genome feature tracks for *M. capitata* and *P. acuta* were derived directly from the published genomes for use in DNA methylation analyses. The *M. capitata* genome annotation yielded gene (a combination of AUGUSTUS and GeMoMa predictions), coding sequence, and intron tracks (Shumaker et al. 2019). Similarly, gene (AUGUSTUS predictions), coding sequence, and intron information was obtained from the *P. acuta* genome (Vidal-Dupiol et al. 2019). Flanking regions 1000 bp upstream and downstream of annotated genes were generated with BEDtools v2.29.2 (`flankBED`) for each genome separately (Quinlan and Hall 2010). Overlaps between genes and flanks were removed from up- or down-stream flanking region tracks using `subtractBED`. Similarly, an intergenic region track was created by finding the complement of genes with `complementBED`, then removing any overlaps with flanking regions using `subtractBED`. All tracks were verified with the Integrative Genomics Viewer (Thorvaldsdóttir, Robinson, and Mesirov 2013). Feature track files generated for both species are available in the project large file repository (Putnam, Roberts, and Venkataraman 2020).

### MBD Enrichment

Before enrichment, DNA (1 µg) in 80 µL Tris HCl (pH 8.0) was sheared to 500 bp using a QSonica Q800R3. Samples were sonicated for 90 sec, with 15 sec on and 15 sec off intervals at 25% amplitude. Fragment length was checked using a D5000 TapeStation System (Agilent Technologies) and samples were sonicated an extra 15 sec to shear DNA from 600 bp to 500 bp as needed.

The MethylMiner kit (Invitrogen; Cat. #ME10025) was used to enrich for methylated DNA prior to MBDBS library generation, with 1µg of input DNA. Manufacturer’s instructions were adhered to with the following modifications: The capture reaction containing the fragmented DNA and MBD beads was incubated with mixing at 4°C overnight, and enriched DNA was obtained with a single fraction elution using 2 M NaCl. Following ethanol addition, samples were centrifuged at 14,000 RCF at 1°C for five minutes. Pellets were resuspended in 25 µL ultra-pure water. Captured DNA was quantified using a Qubit dsDNA HS Kit (Invitrogen).

### MBDBS and WGBS Library Preparation

WGBS and MBDBS libraries were prepared using the Pico Methyl-Seq Library Prep Kit (ZymoResearch Cat. # D5456). Manufacturer’s instructions were followed with the following modifications: For each sample, 1 ng of coral DNA and 0.05 ng of *E. coli* Non-Methylated Genomic DNA (ZymoResearch Cat. # D5016) were used. Samples were always centrifuged at 12,000 RCF for 30 sec with the exception of a 90 sec centrifugation at 12,000 RCFafter the second 200 µL addition of M Wash Buffer. Warmed elution buffer (56°C) was added to each sample to increase DNA elution yield. During the second amplification cycle, 0.5 µL of PreAmp Polymerase was added. After initial clean-up with the DNA Clean and Concentrator kit (ZymoResearch Cat. # D4013), the first amplification step was run for eight cycles. For amplification with i5 and i7 index primers, 1 µL of each primer (10 µM) was used to improve amplification. The volume of the 2X LibraryAmp Master Mix was increased to 14 µL to match the increase in index primer volume.

To remove excess primers from WGBS and MBDBS preparations, samples were cleaned with 11 µL of KAPA pure beads (1X) (KAPA Cat # KK8000) and 80% ethanol. Cleaned samples were resuspended in 12 µL of room-temperature DNA elution buffer from the Pico Methyl-Seq Library Prep Kit. Samples were re-amplified with either two or four cycles, depending on DNA concentration. Re-amplification was conducted with only 0.5 µL of each i5 and i7 index primer (10 µM). After re-amplification, 26 µL of KAPA pure beads (1X) and 80% ethanol were used for clean-up. Final samples were resuspended in 14 µL of room-temperature elution buffer. Primer removal and library size were confirmed by running samples on a D5000 TapeStation System.

### RRBS Library Prep

RRBS libraries were prepared with the EZ DNA RRBS Library Prep Kit (ZymoResearch Cat. # D5460). Manufacturer’s instructions were used with the following modifications: For MspI digestion, 300 ng of input DNA and 15 ng of *E. coli* Non-Methylated Genomic DNA spike were used. Digestions were carried out at 37°C for 4 hours. Adapter ligation was performed overnight, with samples held at 4°C once cycling was completed. Similar to WGBS and MBDBS library preparation, samples were always centrifuged at 12,000 RCF for 30 sec, with the exception of a 90 sec centrifugation after the second 200 µL addition of M Wash Buffer. Warmed elution buffer (65°C) was added to each sample to increase DNA elution yield. Adapter sequences were added via PCR using Index primers following the recommended thermocycling protocol with eleven cycles. Samples were cleaned using 50 µL of KAPA pure beads (1X) and 80% ethanol, then resuspended in 16 µL of the elution buffer. Primer removal and library size were confirmed by running samples on a D5000 TapeStation System.

### DNA Sequence Alignment

All libraries (n = 18) were pooled in equimolar amounts and loaded at 250 pM onto a single Illumina NovaSeq S4 flow cell lane for 2×150 bp sequencing at Genewiz (South Plainfield, NJ). This was estimated to yield 111-138 M reads per library and 99-123x coverage of the *P. acuta* genome (3.3 M bp) and 38-47x coverage of the *M. capitata* genome (8.8 M bp), assuming 100% even coverage (e.g., 150 bp read * 2 pairs * 111 M reads/336,684,533 bp for *P. acuta*).

Sequence quality was checked by FastQC v0.11.8 and adapters from paired-end sequences were trimmed using TrimGalore! version 0.4.5 (Krueger 2012). Following recommendations for methylation sequence analysis from the manufacturer’s protocol and from the Bismark User Guide, 10 bp were hard trimmed from the 5’ and 3’ end of each read for WGBS and MBDBS samples, and RRBS samples were trimmed with `--non_directional` and `--rrbs` options. Bisulfite-converted genomes were created in-silico with Bowtie 2-2.3.4 [Linux x84_64 version; (Langmead and Salzberg 2012)) using `bismark_genome_preparation` through Bismark v0.21.0 (Krueger and Andrews 2011). Trimmed reads were aligned to the BS-converted *P. acuta* genome (Vidal-Dupiol et al. 2019) and the BS-converted *M. capitata* genome (Shumaker et al. 2019) with Bismark v0.21.0 with alignment stringency set by `-score_min L,0,-0.6` and the default MAPQ score threshold of 20. To check mapping rates for endosymbionts and quantify percent methylation, trimmed reads from *P. actua* libraries were also aligned to the *Cladicopium goreaui* genome [type C1, previously *Symbiodinium goreaui* (Liu et al. 2018)] using the same settings as specified above. Reads that mapped ambiguously were excluded and alignment files containing uniquely mapped reads were deduplicated with `deduplicate_bismark` for WGBS and MBDBS samples only. Methylation calls were extracted from sorted deduplicated alignment files using `bismark_methylation_extractor`. Cytosine coverage reports were generated using `coverage2cytosine` with the `--merge_CpG` option to combine methylation data from both strands. Resulting files include bedgraphs and Bismark coverage files (Putnam, Roberts, and Venkataraman 2020). MultiQC v1.8 (Ewels et al. 2016) was run on the trimmed reads, FastQC output, and Bismark reports to assess quality and summarize results.

### Bisulfite conversion efficiency assessment

Trimmed sequence reads were aligned to the genome of *E. coli* strain K-12 MG1655 (Riley et al. 2006) using Bismark v0.21.0 with the `–non_directional` option and alignment stringency set by `-score_min L,0,-0.6`. Bisulfite conversion efficiency was also estimated from coral alignments as the ratio of the sum of unmethylated cytosines in CHG and CHH context to the sum of methylated and unmethylated cytosines in CHG and CHH. Analysis of Variance (ANOVA) was used to test for an effect of library preparation method on conversion efficiency within each species (conversion efficiency ∼ library preparation method) for both coral data estimated and *E. coli* alignment calculated conversion efficiencies. A two-sample t-test was used to test if conversion efficiency calculated from *E. coli* alignments was the same as estimated conversion efficiency for each library preparation method within each coral species.

### Genome-Wide Methylation

General *M. capitata* and *P. acuta* methylation was characterized to describe species-specific patterns. This was carried out by combining BEDgraphs derived from all methods for each species using `unionBedGraphs`. Percent methylation for every CpG locus with at least 5x coverage was averaged, irrespective of how many samples had coverage for that locus. Loci with no data within a method were excluded from downstream analysis. CpGs were classified as being either highly methylated (≥ 50% methylation), moderately methylated (>10% and <50%), or lowly methylated (≤10% methylation).

### Percent Methylation of Shared CpG Loci

Comparisons of percent DNA methylation at CpG loci analyzed by more than one method were performed using the R package `methylKit` (Akalin et al. 2012). A minimum of 5x coverage was required across all samples for a CpG locus to be considered in the analyses. The `unite` function in `methylKit` was used to identify CpG loci that were covered across all 9 samples (3 individuals per method) per species. Scatterplots and Pearson correlation coefficients were calculated using the function `getCorrelation`. Additionally, differential methylation tests were performed on pairwise comparisons between methods (WGBS versus RRBS, WGBS versus MBDBS, and RRBS versus MBDBS). Discordant methylation was quantified using a logistic regression model on CpG loci that were covered across all 6 samples (3 samples from each method compared) in each pairwise comparison using the `calculateDiffMeth` function with default parameters.

### CpG Coverage

To assess average genome-wide CpG coverage, the number of cytosines passing different read depth thresholds (5x, 10x, 15x, 20x, 25x, 30x, 40x, and 50x) were totaled from the CpG coverage reports output by the Bismark `coverage2cytosine` function (detailed above) for each sample. These totaled CpGs were then relativized to the number of CpGs in their respective genomes (*M. capitata,* 28,684,519 CpGs; *P. acuta,* 9,155,620 CpGs). Next, average and standard deviation of genome-wide CpG fractions were calculated for each method within each species (n = 3), and these were plotted across different read depth thresholds using `ggplot2` (Gómez-Rubio 2017).

To estimate overall genome-wide CpG coverage, a downsampling analysis was performed by pooling all sample reads within a method and species. Briefly, trimmed fastq files were concatenated for each method and species, then randomly subsampled to 50, 100, 150, and 200 million reads. Next, alignment and methylation calling were carried out as described above on each subset, and the number of cytosines with 5 or more reads were totaled from CpG coverage reports from each subset. Sequencing saturation was estimated from a Michaelis-Menten model with the `drm` function from the R package `drc` (Ritz et al. 2015) using CpG coverage reports from subsampled data as input. Both observed CpG coverage from subsampled data and estimated CpG coverage were plotted using the R package `ggplot2` (Gómez-Rubio, 2017).

### Gene Coverage

To compare the differences in genes with methylation data by method, we identified the CpGs with 5x coverage that intersected with gene regions using `intersectBED` (bedtools v2.30.0). The proportion of genes with methylation data was calculated by identifying genes in each method that had 5x CpG data and dividing by the total number of genes in the reference genome. We assessed library preparation method bias on functional information by considering gene ontology (GO) terms associated with genes containing CpG data (at least one CpG per gene with 5x coverage in any library). For the set of genes with CpG coverage we performed enrichment analysis to determine if these genes resulted in significant enrichment of particular GO terms using GOseq (Young et al. 2010) and accounting for gene length.

### Proportion of Detected CpGs for Orthologs

To describe the differences in DNA methylation detected by each method at a more functional level, and given the connection of gene body methylation and gene expression in invertebrates (Roberts and Gavery 2012) and corals specifically (Liew et al. 2018), the presence of CpG data within all genes was calculated for each species, by method. First, a CpG gff track was generated using EMBOSS (Rice, Bleasby, and Ison 2011) with the `fuzznuc` command searching for the pattern CG. For each sample, `intersectBED` was used to identify CpGs with 5x coverage that intersected with gene regions. This was also done for the reference genome CpG gff track. CpG counts per gene were compiled for each sample and the mean taken per method. The proportion of CpGs per orthologous gene was calculated by dividing the mean number of CpGs with 5x coverage from the three samples per method and dividing that by the number of CpG possible summed per gene from the reference genome CpG gff track. The proportion of CpG data in a gene was then visualized in heatmaps for all genes of *M. capitata* and *P. acuta*.

### Genomic Location of CpGs

For both *M. capitata* and *P. acuta*, the overlap between genome feature tracks and species-specific CpG data at 5x coverage was characterized with BEDtools v2.29.2 to assess the presence CpGs in various regions by method (Quinlan and Hall 2010). Since only gene, coding sequence, intron, flanking regions, and intergenic region tracks were common between species, these were the tracks used in downstream analyses. A combination of PCoA, PERMANOVA and beta-dispersion tests, and chi-squared contingency tests were used to determine if the library preparation method influenced the proportion of CpGs detected in a specific genomic feature. A separate contingency test was used for each genomic feature.

Code for all calculations can be found in (Putnam, Roberts, and Venkataraman 2020)

## Results

To compare the performance of bisulfite sequencing methods in the reef-building scleractinian corals *Montipora capitata* and *Pocillopora acuta*, we isolated DNA and generated WGBS, RRBS, and MBDBS libraries for three individuals from each species to yield a total of 18 libraries (**Figure 1**).

### DNA Sequence Alignment

Sequencing of all 18 libraries resulted in 1.82 x 10^9^ read pairs, of which 99.1% remained after QC and trimming (Additional file 1: **<u>Table ST1</u>**). Individual libraries were generally sequenced to the same depth (8.1 ± 0.34 x 10^7^ reads; mean ± S.E.) across library preparation methods and species, with the exception of *P. acuta* RRBS libraries, which were sequenced 2- to 4-fold deeper (19.2 ± 4.67 x 10^7^ reads; mean ± S.E.). The average mapping efficiencies for all *P. acuta* and all *M. capitata* libraries were 45.5 ± 5.2% and 38.9 ± 4.3% of reads, respectively (Additional file 2: **<u>Table ST2</u>**). In comparison to other methods, MBDBS libraries had a larger proportion of reads (73.1% ± 9.9%) that did not align to the coral genomes (Additional file 3: **<u>Figure SF1</u>**). To investigate this we aligned *P. acuta* libraries to a known symbiont *Cladocopium goreaui* genome (C1 (Liu et al. 2018)) for which the genome sequence was available at the time of analysis. We found a sizable proportion of the MBDBS reads mapped to the symbiont genome (23.6 ± 10.6%), while a much smaller proportion of RRBS and WGBS reads mapped to the symbiont genome (5.04 ± 0.22% and 1.92 ± 0.3% respectively) (Additional file 4: **<u>Table ST3</u>**).

### Bisulfite conversion efficiency assessment

Bisulfite conversion efficiency calculated from alignments of the *E. coli* Non-Methylated Genomic DNA spike-in ranged from 98.6 to 99.3% in *M. capitata* and from 98.3 to 99.1% in *P. acuta* (Additional file 5: **<u>Table ST4</u>**), and this differed by library preparation method for both *M. capitata* (F_2,6_= 114.22, *P* = 1.676×10^-05^) and *P. acuta* (F_2,6_= 7.24, *P* = 0.025) libraries. In general, conversion efficiency calculated from the *E. coli* alignments did not differ from conversion efficiency estimates from CHG and CHH methylation (under the assumption that non-CpG methylation does not occur in corals, see also (Liew et al. 2018)) from coral alignments in *M. capitata* and *P. acuta*. (Additional file 6: **<u>Figure SF2</u>** <u>and Additional file 7: **Table ST5**</u>).

### Methylation characterization

For each species, the general methylation landscape was characterized for CpG loci with 5x coverage identified in any method. The *M. capitata* genome was more methylated than *P. acuta* (**Figure 2**). Using a cutoff of ≥ 50% methylation to define methylated CpGs, of the 13,340,268 CpGs covered by the *M. capitata* data, 11.4% were methylated. In contrast, only 2.9% of the 7,326,297 CpGs in *P. acuta* were methylated.

**Figure 2.**
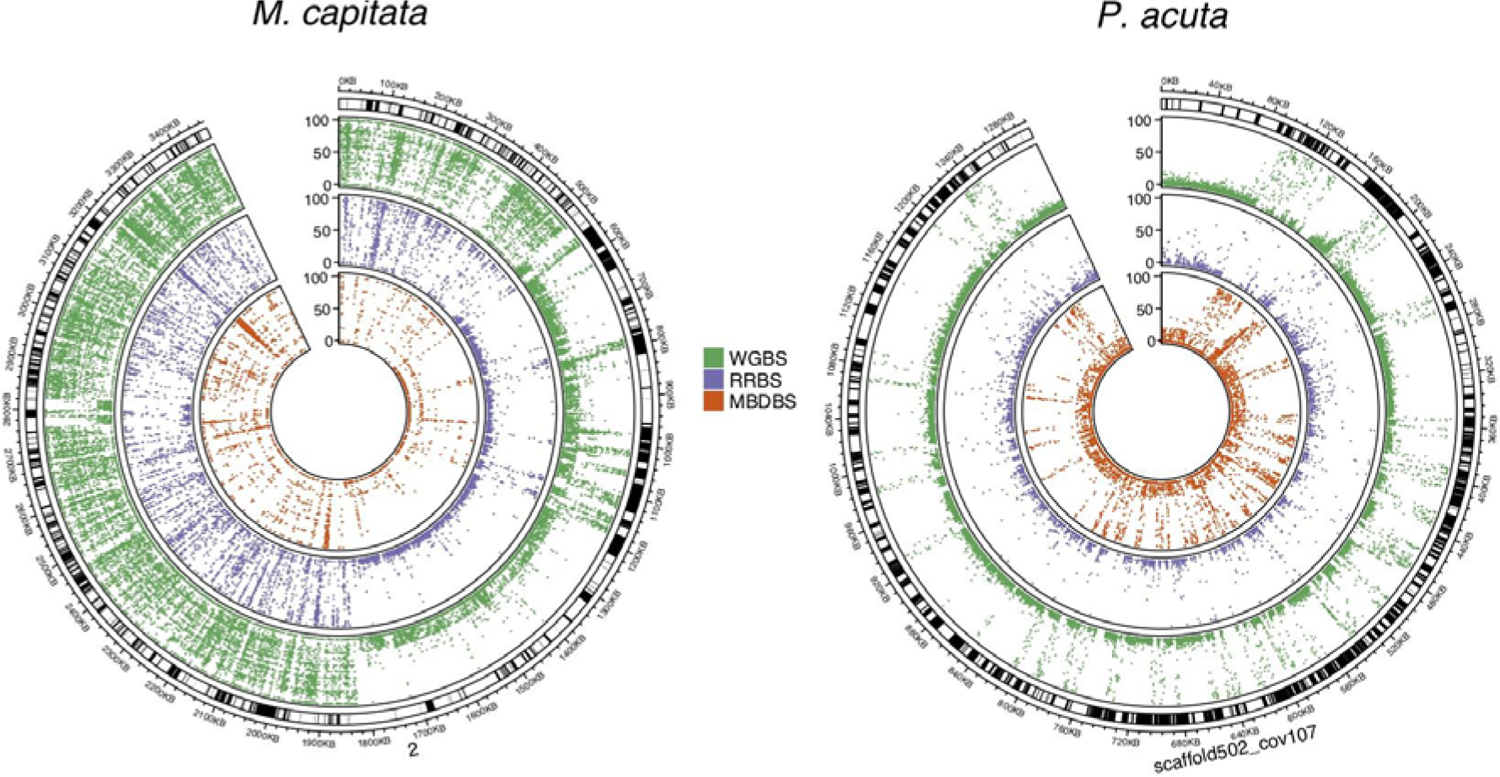
Mean percent methylation of CpGs. Data is presented for CpGs with 5x coverage for each method on the largest scaffolds of each genome. The outer track shows the scaffold locations and dots indicate the percent methylation as indicated by the y-axes from 0-100% for each of the inner tracks.

Both genomes were predominantly lowly methylated (≤ 10% methylated): 79.6% CpGs in *M. capitata* and 91.3% CpGs in *P. acuta* were lowly methylated. The remaining 9.0% of CpGs in *M. capitata* and 5.8% of CpGs in *P. acuta* were moderately methylated (10-50% methylation). The different methods captured varying proportions of highly, moderately, and lowly methylated CpGs (Additional file 8: **<u>Figure SF3</u>**).

### Correlation of methylation among common CpG loci

For quantitative comparison of method performance, we reduced the dataset to loci covered at 5x read depth across all methods and samples for each species, referred to here as ‘shared loci’. The number of shared loci was 4,666 CpG for *M. capitata* and 93,714 CpG for *P. acuta*. A PCA of CpG methylation for loci covered at 5x read depth showed that libraries tended to cluster in PC space by preparation method, rather than by individual (Additional file 9: **<u>Figure SF4</u>**). Variation in methylation levels of the shared loci across all *M. capitata* samples was lower within a method than between methods (Additional file 9: **<u>Figure SF4A</u>**). For *P. acuta*, RRBS and WGBS methods showed similar methylation levels of shared loci, but these were different from the methylation level of loci identified in MBDBS (Additional file 9: **<u>Figure SF4B</u>**). To further explore the variation in methylation observed by method, we directly correlated quantitative methylation calls for the shared loci (**Figure 3**). Correlations among biological replicates within a method were higher and less variable for *M. capitata* compared to *P. acuta*. Correlations between pairs of methods for *M. capitata* ranged on average from 0.75-0.82, whereas correlations for *P. acuta* ranged from 0.40-0.64. For *M. capitata*, WGBS versus MBDBS had the highest correlation. For *P. acuta*, WGBS versus RRBS had the highest correlation.

**Figure 3.**
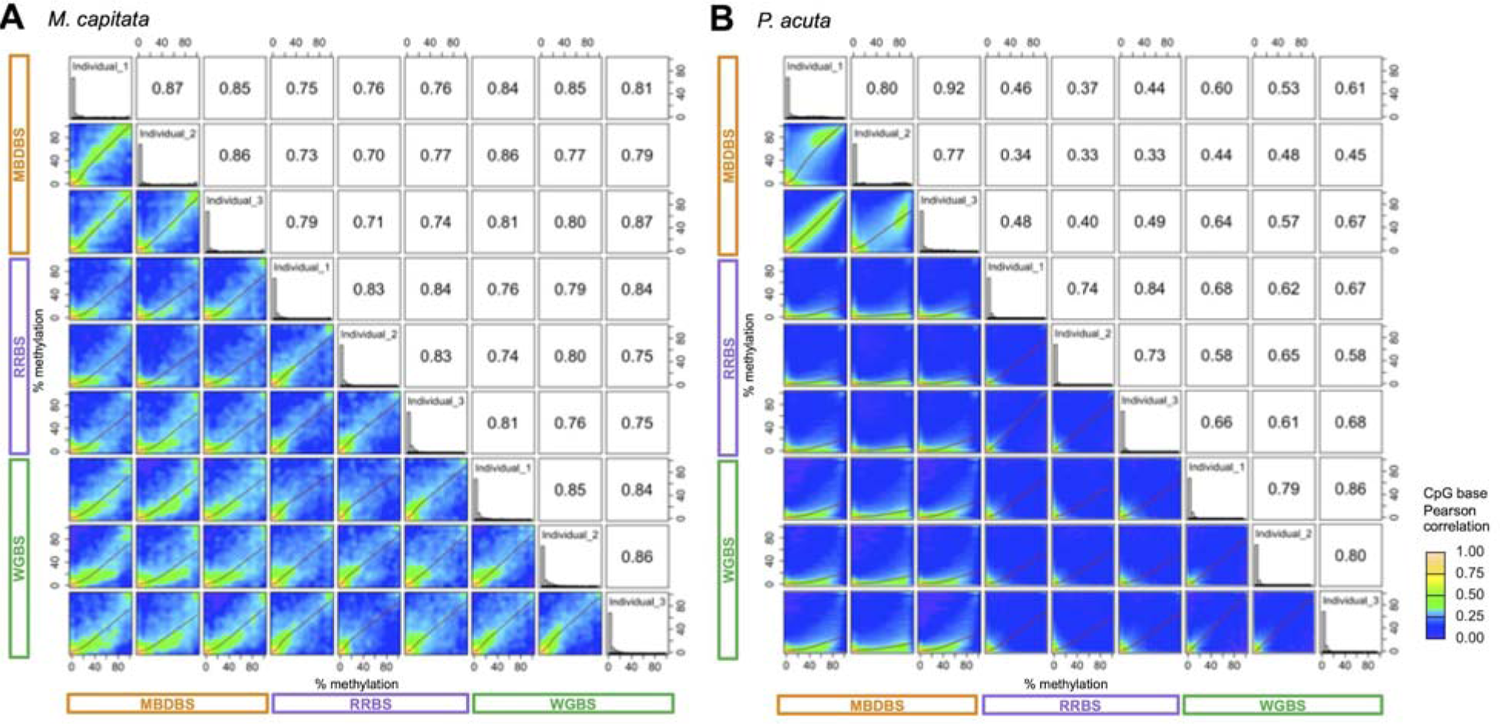
Matrix of pairwise scatter plots for shared CpG loci. Data is presented for CpG covered at ≥ 5x across all samples) for A) *M. capitata* (n=4,666 common loci) and B) *P. acuta* (n=93,714 common loci). The red lines represent linear regression fits and the green lines are polynomial regression fits. Pearson correlation coefficients for each pairwise comparison are presented in the upper right boxes. Methods are color coded on the X and Y axes (WGBS = green, MBDBS = purple, and RRBS = orange) and replicate samples are indicated on the diagonal along with histograms of % CpG methylation.

Discordance in methylation quantification between methods was evaluated by identifying the number of CpG loci with large differences (>50%) in methylation for each species. WGBS versus RRBS showed the lowest discordance in both species (0.4% for *M. capitata* and 0.5% for *P. acuta*). The highest discordance in methylation was found in comparisons with MBDBS for *P. acuta*, with 11% and 15% of CpG sites being called at least 50% different for comparisons with WGBS and RRBS, respectively. In contrast, only 0.4% and 5% of common CpG sites were at least 50% different between MBDBS versus WGBS and MBDBS versus RRBS, respectively, for *M. capitata*. A majority of the discordance was due to higher methylation calls in MBDBS compared to WGBS or RRBS (**Figure 3B**).

### CpG coverage

Consistent with what would be expected based on genome size, *P. acuta* libraries have higher genome-wide CpG coverage than *M. capitata* regardless of library preparation method (**Figure 4A-C**). For both species, WGBS and MBDBS libraries covered more CpGs than RRBS libraries, whereas RRBS libraries tended to show greater read depth for the CpGs that it did cover. In other words, at >20x read depth, RRBS libraries covered more CpGs than either WGBS or MBDBS (**Figure 4** insets). Modelling increased sequencing depth for RRBS or MBDBS libraries showed little impact on the fraction of genome-wide CpGs covered in *M. capitata*, while increasing sequencing depth from 50 M to 200 M for WGBS libraries in both species and for MBDBS in *P. acuta* showed a substantially larger fraction of CpGs covered (Additional file 10: **<u>Figure SF5</u>**).

**Figure 4.**
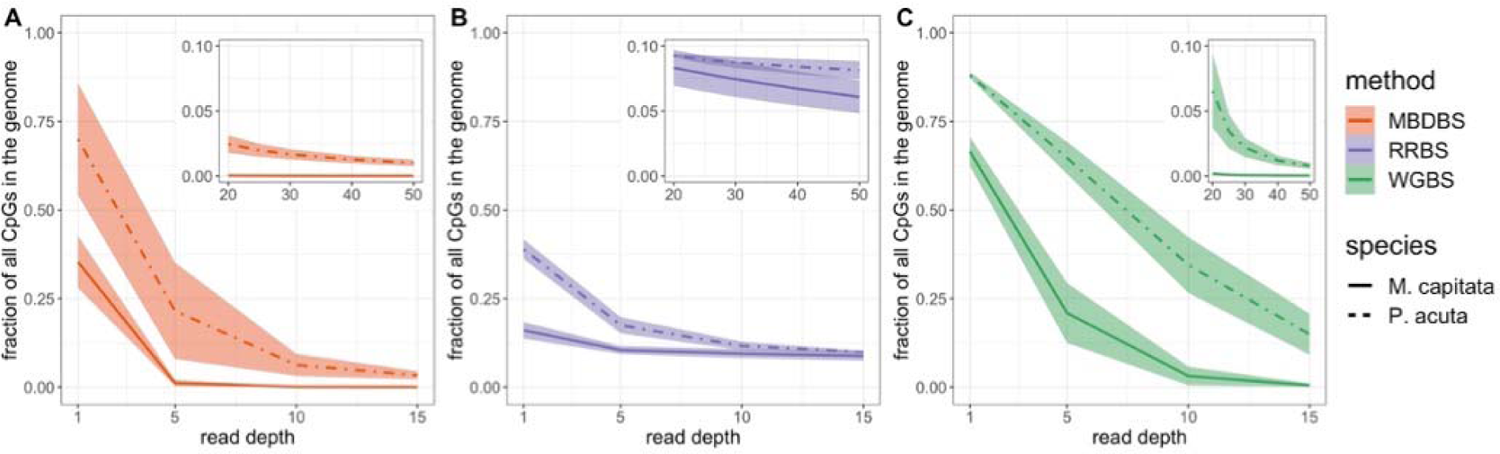
CpG site coverage across library preparation methods. Mean fraction of CpG sites in the genome covered at different sequencing depths (read depths) by (**A**) MBDBS libraries, (**B**) RRBS libraries, and (**C**) WGBS libraries with standard deviations shown by shaded areas (see Additional file 2: **<u>Table ST2</u>** for number of reads in each sample).

### Gene Coverage

For each species, WGBS provided CpG data for the highest proportion of genes in the genome: 84.5% and 94.3% for *M. capitata* and *P. acuta*, respectively. RRBS generated CpG data for 54.7% and 63.5% of the genes for *M. capitata* and *P. acuta*, respectively. MBDBS provided the most divergent coverage of genes with 44.1% and 86.9% for *M. capitata* and *P. acuta*, respectively. When performing functional gene enrichment on the gene sets that contained CpG data for each method, there were multiple enriched GO categories for all methods and both species (**Figures <u>SF6</u>, <u>SF7</u>)**. For *M. capitata*, there were 223, 401, and 253 enriched molecular function (MF) terms (**<u>Table ST6</u>**) and 508, 1123, and 853 biological process (BP) terms for WGBS, RRBS, and MBDBS, respectively (**<u>Table ST7</u>**). For *P. acuta* there were 164, 282, and 287 terms enriched for MF (**<u>Table ST8</u>**) and 313, 749, and 682 terms enriched for BP for WGBS, RRBS, and MBDBS, respectively, (**<u>Table ST9</u>**).

### CpG coverage within orthologous genes

In order to assess the potential for cross-species comparisons using an equivalent dataset we quantified CpG data available across one-to-one orthologous genes. For *M. capitata*, WGBS yielded the highest proportion of CpGs, followed by RRBS, and then MBDBS (Additional file 11: **Figure <u>SF8A</u>**). This differed in *P. acuta* with WGBS yielding the highest proportion of CpGs on average across orthologs, followed by MBDBS, and then RRBS (Additional file 11: **Figure <u>SF8B</u>**).

### Genomic location of CpGs

In order to compare locations of CpG data between genomic features for each species and method, all CpGs with 5x coverage were characterized based on genomic feature location (**Figure 5**). Global PERMANOVA tests found significant differences between library preparation methods for CpG coverage in various genome features for *M. capitata* and *P. acuta* (Additional file 12: **Table <u>ST10</u>**). Although post-hoc pairwise PERMANOVA tests did not reveal differences between sequencing methods, power for these was probably low (due to low sample size). Pairwise chi-squared tests indicated there are differences in CpG location for both species. In particular, CpGs in gene bodies were significantly enriched over other genomic features with MBDBS (**Figure 5**; Additional file 13: **Table <u>ST11</u>**). Visual inspection of PCoA also revealed the proportion of CpGs captured in coding sequences (CDS) drove differences between MBDBS and the other methods in both species (**Figure 5C-D**).

**Figure 5.**
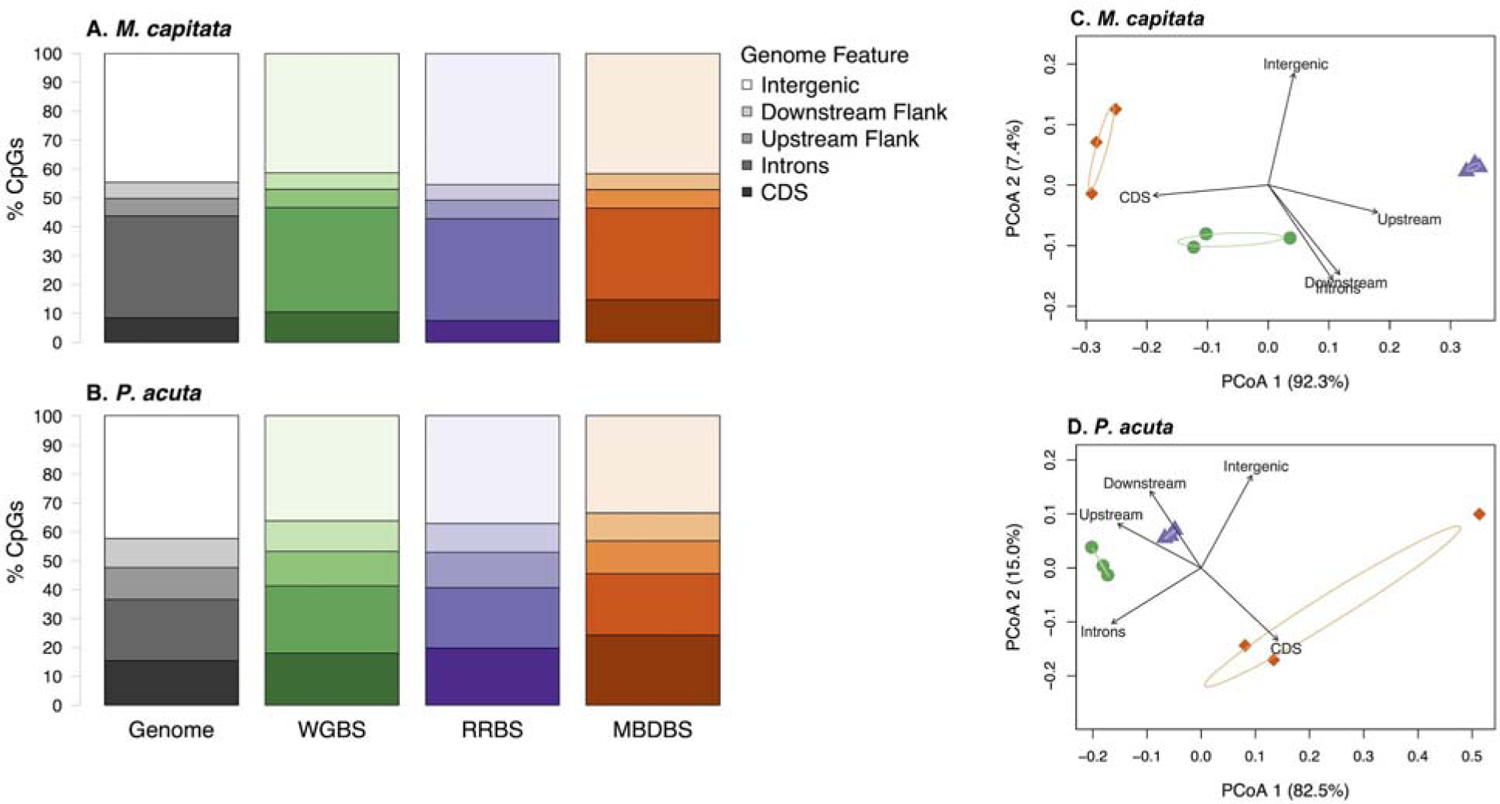
Percent of CpGs detected by sequencing methods in genome features. A) for *M. capitata* and B) *P. acuta*. Genome features considered were coding sequences (CDS), introns, 1 Kb flanking regions upstream (Upstream Flank) or downstream of genes (Downstream Flank), and intergenic regions. Each bar corresponds to all the CpGs in the genome (Genome), or each method (WGBS, RRBS, or MBDBS). The order of the genome features depicted in each bar is identical to what is displayed in the legend, with the darkest shade representing CDS, and the lightest shade representing intergenic regions. Principal Coordinate Analyses associated with PERMANOVA and beta-dispersion tests related to Additional file 12: **<u>Table ST6</u>** that show differences in proportion of CpGs in various genomic locations (CDS, introns, upstream flanks, downstream flanks, and intergenic regions) for **C**) M. capitata and **D**) P. acuta. WGBS is represented by green circles, RRBS by purple triangles, and MBDBS by orange diamonds. Percent variation explained by each PCoA axis is included in the axis label. Ellipses depict 95% confidence intervals for each sequencing method. All eigenvectors are significant at the α = 0.05 level.

## Discussion

We evaluated the performance of three approaches that use bisulfite-treated DNA for library preparation to enable single base-pair resolution quantification of DNA methylation in corals. Our results demonstrate that the methylation landscape can vary significantly across species, which is a critical consideration for both interpreting environmental response capacity, and therefore for experimental design. Whereas WGBS is the gold standard for studying methylation, it comes at a high cost. MBDBS enriches for gene regions, which may be useful for taxa with gene body methylation. On the other hand, RRBS provides greater coverage depth for a smaller fraction of the genome, but lacks specificity for genomic features or DNA methylation. Taken together, our findings indicate biology, genome architecture, regions of interest, and depth of coverage are critical considerations when choosing methods for high resolution quantification of DNA methylation profiles in invertebrates.

*M. capitata* has a relatively high environmental tolerance (Bahr, Jokiel, and Rodgers 2016; Gibbin et al. 2015; Putnam, Davidson, and Gates 2016), which has previously been attributed to its symbiont composition (Cunning, Ritson-Williams, and Gates 2016), genome characteristics (Shumaker et al. 2019), perforate tissue-skeletal architecture and tissue thickness, and heterotrophic capacity (Rodrigues and Grottoli 2007). Of particular relevance to DNA methylation are genomic aspects such as gene family duplication and high repeat content in *M. capitata* (Shumaker et al. 2019). We found overall DNA methylation was higher in *M. capitata* than in *P. acuta*, supporting early bulk analyses of DNA methylation in these species (Putnam, Davidson, and Gates 2016). While the predicted number of genes is similar, the genome size of *M. capitata* is over twice that of *P. acuta* (Shumaker et al. 2019; Vidal-Dupiol et al. 2019). One explanation for the higher methylation in *M. capitata* is that with greater energy availability — through translocation from high density Symbiodiniaceae populations and energy stores in perforate tissues — there is greater capacity for maintenance methyltransferase to maintain high methylation, and thus reduce gene expression variability and spurious expression (Li et al. 2018; Liew et al. 2018). High constitutive methylation could allow “frontloading” of stress response genes (e.g., (Barshis et al. 2013)), providing greater stress tolerance. Another possible explanation is that the higher level of methylation contributes to the silencing of repeated genetic elements. In contrast, with a small and non-repetitive genome, imperforate thin tissues, and low energy reserves, *P. acut*a may be more energetically limited. Thus, *P. acuta* may be expected to show lower DNA methylation across the genome as we demonstrate here, as well as a higher propensity for inducible methylation in the presence of stressors (Putnam, Davidson, and Gates 2016).

Another striking contrast in DNA methylation in these species is the lack of concordance in the percent methylation values for *P. acuta* among methods compared to *M. capitata* (**Figure 3**). The potential for chimerism in corals (Schweinsberg et al. 2015; Oury, Gélin, and Magalon 2020) and differences in tissue structure (e.g., perforate or imperforate) between species could contribute to differences in concordance across methods for quantifying DNA methylation. One possibility is that Pocilloporids are chimeric and multiple genotypes exist (Schweinsberg et al. 2015; Oury, Gélin, and Magalon 2020). Although percent DNA methylation concordance across methods was generally high, in *P. acuta* there was approximately a 10% higher level of discordance in percent methylation quantification when comparing WGBS to RRBS or MBDBS (**Figure 3**). This discordance could have resulted from differences in *P. acuta* and *M. capitata* tissue structure. There is the potential to homogenize and extract DNA from all cell types from the thin, imperforate tissues of *P. acuta*, as opposed to the thick, perforate tissues in *M. capitata* (Putnam et al. 2017), likely contributing to a greater number of cell types, and thus methylation differences, captured in our *P. acuta* samples. Furthermore, the microhabitats created in the tissues of these two species likely differ substantially spatially (Putnam et al. 2017), creating cell-to-cell variability in methylation content. Since the likelihood of capturing multiple cell types in bulk DNA extractions varies with tissue structure, future studies should consider methods such as fluorescence-activated cell sorting (Rosental et al. 2017; Hu et al. 2020), or laser microdissection (e.g., (Liew et al. 2018)), to target specific tissues or cell types and reduce cell-to-cell methylation variability. Whereas this does not necessarily indicate a bias in our methods, it highlights the need to account for the biological characteristics of a species when designing an experiment and evaluating results. Also when comparing across species, given genetic-epigenetic correlations, particularly in the case of DNA methylation and the requirement for a CpG sequence target site (Dimond and Roberts 2020; Johnson et al. 2020), variation in genome architecture, gene number, and content will impact the presence and use of DNA methylation as a mechanism of gene expression regulation.

The gold standard for bisulfite sequencing, WGBS, can be cost prohibitive particularly if comparing multiple species and treatments. As expected, we found that WGBS performed well, particularly for *P. acuta* which has a smaller genome. We found gene ontology enrichment of the genes covered by CpG data was affected by library preparation method, likely attributed to genome characteristics and relatively higher methylation in *M. capitata* affecting the genes covered by reduced representation methods (e.g. 44.1% of genes in *M. capitata* and 86.9% of genes in *P. acuta* covered by MBDBS). There may be preferential enrichment of hypermethylated genes in *M. capitata* by MBD and thus MBD in *M. capitata* may not capture the breadth of genes found in the less methylated *P. acuta*. Therefore MBDBS approaches could benefit from species-specific protocol optimization (Aberg et al. 2018). GO enrichment analysis identified changes in the significantly enriched GO terms as the proportion of genes with data decreases (Figure SF7 and SF8) and these GO terms were found across many BP and MF terms, not limited to a particular set (Tables ST6-ST9). Focusing on direct comparison of gene orthologs, WGBS performed the best in terms of data for CpGs per gene. Based on the gene ortholog comparisons, MBDBS provided more information than RRBS for *P. acuta*, however the opposite held true for *M. capitata.* Collectively, these differences are likely attributable to the different genome size, assembly quality, and/or inherent differences in methylation that result in differential enrichment.

For both species, WGBS and MBDBS libraries covered more CpGs than RRBS libraries; however, RRBS libraries showed greater read depth for CpGs. This is because RRBS subsampled a specific, smaller portion of the genome than MBDBS or WGBS, allowing more read coverage. Hence, CpG coverage did not largely increase when deeper sequencing was modeled using RRBS data (Additional file 10: **<u>Figure SF5</u>**). RRBS was designed to enrich for CpG islands, short stretches of DNA with higher levels of CpGs (∼1 CpG per 10bp), that are typically found in mammalian promoters and enhancer regions and thought to play a role in gene regulation (Gu et al. 2011). We found RRBS yielded a well-covered reduced representation of the genome, which is important for bisulfite data where high read depth is desired, and locus methylation levels were concordant with WGBS for both species. However, RRBS did not enrich for promoters or other particular genomic regions compared to the other bisulfite sequencing methods (**Figure 5**), and in fact tended to identify unmethylated regions. For this reason, RRBS is not the best choice for gene-focused methylation studies in corals and other invertebrate taxa with gene body methylation.

A critical consideration in deciding to perform MBDBS in corals is the amount of DNA methylation present from any symbiont. If any non-target organisms have substantially more DNA methylation than the target organism, MBDBS data could become saturated by methylated DNA from non-target organisms, lowering sampling of the target species. We observed this in *P. acuta*, for which we had the genome of its Symbiodiniaceae which has ∼90% genome-wide methylation (Lohuis and Miller 1998; de Mendoza et al. 2018). When compared to RRBS and WGBS data, we found a 4 to 10-fold enrichment of Symbiodiniaceae DNA in *P. acuta* MBDBS data. Separation of host and symbionts is therefore recommended to obtain the greatest read counts for the organism of interest, but this comes at the cost of not being able to obtain RNA from the same nucleic acid pool. For example, physical separation of the host and symbiont in living cells impacts gene expression and attempts at physical separation after freezing can degrade the host RNA. Simultaneous extraction of holobiont RNA and DNA from the same nucleic acid pool provides the optimal approach for detecting interactions between DNA methylation and epigenetic regulation of gene expression. This comes at the cost of generating excess reads to overcome highly methylated Symbiodiniaceae DNA.

MBDBS can enrich for gene regions in species where methylation is primarily found in gene bodies such as in corals (reviewed in (Eirin-Lopez and Putnam 2019)), and can thus provide insight into mechanisms underlying physiological or organismal responses. We found that MBDBS significantly enriched for gene bodies, specifically CDS and introns, when compared to RRBS and WGBS in both *M. capitata* and *P. acuta* (**Figure 5**). While MBDBS may be a good choice to examine gene body methylation at a reduced cost, species differences in CpG coverage within orthologous genes with MBDBS (Additional file 10: **<u>Figure SF</u>6**) may complicate cross-species comparisons by reducing the amount of data available for analysis. Additionally, we found high discordance between MBDBS and non-enrichment methods, WGBS and RRBS, for *P. acuta*. MBDBS is the only method we evaluated that can non-randomly sub-sample genomes present in a DNA sample through preferential pull-down of methylated DNA. Differences in methylation across the sampled genomes could result from cell-to-cell heterogeneity in methylation or cell-type (e.g., methylation of calcifying cells may differ from symbiont hosting cells). In other words, MBDBS data may represent only a subpopulation of highly methylated cells, while WGBS and RRBS represent the average methylation across all cells in the sample. Using a consistent tissue type is important to limit potential methylation heterogeneity, and caution should be taken when comparing MBDBS data directly to that of non-enrichment bisulfite sequencing approaches.

Although MBDBS did enrich for methylated regions of the genome, 80% of CpGs in *M. capitata* and 82% of CpGs in *P. acuta* interrogated with MBDBS were lowly methylated (<10% methylated) (Additional file 8: **<u>Figure SF3</u>**). This is expected and is consistent with previous reports applying MBDBS in other marine invertebrates where unmethylated CpGs actually represent the highest proportion of loci in the data, attributable to the nature of the methylation landscape and enrichment protocol (e.g. (Gavery and Roberts 2013; Venkataraman et al. 2020)). The base-pair resolution of methylation revealed by MBDBS is a benefit over MBD-Seq alone because it enables a fine-scale examination of specific genomic features (e.g. exon-intron boundaries) that may not be possible with the regional resolution of MBD-Seq. Without complete knowledge of the relative importance of a single loci compared to a region, it is difficult to compare trade-offs between MBDBS and MBD-seq. However, bisulfite sequencing requires significant coverage to quantify DNA methylation.

MBDBS may have potential biases that should be considered when interpreting results. If a treatment comparison, population comparison, or developmental change results in a given region (∼500bp) going from being highly methylated to fully unmethylated, then it is likely that this region would not be interrogated by MBDBS, due to an absence of data in the unmethylated condition. This is a potential source of bias in MBDBS data and may contribute to important differentially methylated regions being overlooked: for example if one treatment results in high methylation and is captured by MBDBS and another treatment results in no methylation and is not captured by MBDBS, this region would be filtered out of the analysis because of missing data in some individuals. Further, the potential of MBDBS to provide limited information for unmethylated genes may introduce bias in studies that seek to draw relationships between methylation level and gene expression. Just as with many interpretations of key findings we present, a more complete understanding of the mechanistic functional role DNA methylation plays in genome regulation in the species of interest is needed.

There is a greater capacity to gain mechanistic insight when using methods that have single base-pair resolution of methylation data compared to methylation enrichment without bisulfite treatment or to bulk percent methylation approaches. For example, hypotheses such as the linkage between DNA methylation and alternative splicing (Roberts and Gavery 2012) are more accurately tested with bisulfite sequencing approaches. We acknowledge the cost of generating genomic resources and bisulfite sequencing data can be higher than other approaches. While WGBS is supported here as the gold standard for DNA methylation quantification, consideration should be given to specific study hypotheses in light of the pros and cons of the enrichment or reduced representation approaches presented here and in other comparative works (Dixon and Matz 2020). Our results suggest that it would be unwise to use multiple different library preparation methods for comparing individuals within a study, especially for studies in which familial relationships are to be compared. As technology advances, it would be ideal to move away from harsh bisulfite conversion to assess DNA methylation with single base-pair resolution across whole genomes in the absence of DNA treatment (e.g., Oxford Nanopore).

Our results provide a quantitative comparative assessment that can be used to inform the choice of sequencing DNA methylation in corals and other non-model invertebrates. Together these metrics enable comparative capacity for three common methods in two coral taxa that vary in their phylogeny, genome size, symbiotic unions, and environmental performance, and thus provide the community with a more comprehensive foundation upon which to build laboratory and statistical analyses of DNA methylation, plasticity, and acclimatization.

## Abbreviations

CpG: cytosine and guanine separated by a phosphate group

ELISA: Enzyme-Linked Immunosorbent Assay

MBD-seq: Methyl-CpG Binding Domain Sequencing

MBDBS: Methyl-CpG Binding Domain Bisulfite Sequencing

RRBS: Reduced representation bisulfite sequencing

RCF: Relative Centrifugal Force

WGBS: Whole genome bisulfite sequencing

## Acknowledgements

We thank Putnam Lab members E. Strand and M. E. Schedl for their technical support. Data analyses were facilitated through the use of advanced computational, storage, and networking infrastructure provided by the Hyak supercomputer system at the University of Washington.

## Data accessibility

The datasets supporting the conclusions of this article are available in the Coral Methylation Methods Comparison repository, http://doi.org/10.17605/OSF.IO/X5WAZ, and included within the article and its additional files. All raw data can be accessed under NCBI Bioproject PRJNA691891.

## Author contributions

HMP designed the study and collected the samples. YRV, SAT, MRG, HMP, and SBR performed data analyses and drafted the manuscript. DB, AD-W, JME-L, KMJ, KEL, and JBP provided analytical insight and manuscript revisions. All authors read and approved the final manuscript.

## Ethics declarations

Ethics approval and consent to participate

The coral samples were collected under Hawai i Department of Aquatic Resources Special Activity Permit SAP 2019-60.

## Consent for publication

Not applicable.

## Supplementary Information

