## Supporting Information for "Invertebrate methylomes provide insight into mechanisms of environmental tolerance and reveal methodological biases"

**Table of Contents:**

[**Table ST1.csv**](https://github.com/hputnam/Meth_Compare/blob/master/output/supplemental-material/ST1.csv)

Trimming statistics.

Descriptive statistics of sequencing reads resulting from sequencing adapter trimming.

[**Table ST2.csv**](https://github.com/hputnam/Meth_Compare/blob/master/output/supplemental-material/ST2.csv)

Alignment statistics.

Descriptive statistics of sequencing reads resulting from Bismark alignment to coral genomes.

[**Table ST3.csv**](https://github.com/hputnam/Meth_Compare/blob/master/output/supplemental-material/ST3.csv)

C1 alignment statistics.

Descriptive statistics of sequencing reads resulting from Bismark alignment to the symbiont *Cladocopium* *goreaui* (C1) genome.

[**Table ST4.csv**](https://github.com/hputnam/Meth_Compare/blob/master/output/supplemental-material/ST4.csv)

*E. coli* unmethylated DNA spike conversion efficiency and alignment statistics.

Descriptive statistics of sequencing reads resulting from Bismark alignment to the *E. coli* genome.

[**Table ST5.csv**](https://github.com/hputnam/Meth_Compare/blob/master/output/supplemental-material/ST5.csv)

Bisulfite conversion efficiency statistics.

Results from ANOVA test comparing conversion efficiency calculated from *E. coli* unmethylated DNA spike alignments and conversion efficiency estimated from non-CpG methylation from coral alignments.

[**Table ST6.csv**](https://github.com/hputnam/Meth_Compare/blob/master/output/supplemental-material/ST6.csv)

Gene ontology enrichment of Molecular Function terms from gene sets with CpG data for the three library prep methods for *M. capitata*.

[**Table ST7.csv**](https://github.com/hputnam/Meth_Compare/blob/master/output/supplemental-material/ST7.csv)

Gene ontology enrichment of Biological Process terms from gene sets with CpG data for the three library prep methods for *M. capitata*.

[**Table ST8.csv**](https://github.com/hputnam/Meth_Compare/blob/master/output/supplemental-material/ST8.csv)

Gene ontology enrichment of Molecular Function terms from gene sets with CpG data for the three library prep methods for *M. capitata*.

[**Table ST9.csv**](https://github.com/hputnam/Meth_Compare/blob/master/output/supplemental-material/ST9.csv)

Gene ontology enrichment of Biological Process terms from gene sets with CpG data for the three library prep methods for *M. capitata*.

[**Table ST10.csv**](https://github.com/hputnam/Meth_Compare/blob/master/output/supplemental-material/ST10.csv)

Results from PERMANOVA and beta-dispersion tests for genomic location

PERMANOVA R^2^ and Adonis *P*-value, and beta dispersion test *F*-statistic and ANOVA *P*-value for obtained by comparing CpG genomic location between methods for *M. capitata* and *P. acuta*.

[**Table ST11.csv**](https://github.com/hputnam/Meth_Compare/blob/master/output/supplemental-material/ST11.csv)

Contingency test results for *M. capitata* and *P. acuta.*

Test statistics and *P*-values from chi-squared tests comparing CpG methylation status and genomic location between WGBS, RRBS, and MBDBS for *M. capitata* and *P. acuta*.

**Figure SF1.jpg**

Alignment information.

Summary of sequencing depth and alignments for all libraries. Bars show average number of reads for each method and species and error bars show standard deviation.

**Figure SF2.jpg**

Bisulfite conversion efficiency assessment.
Bisulfite conversion efficiency calculated from *E. coli* unmethylated DNA spike alignments or estimated from non-CpG methylation from coral alignments for *M. capitata* libraries and *P. acuta* libraries.

**Figure SF3.jpg**

Comparison of CpG methylation status.

Percent of highly methylated (≥ 50%; darkest shade), moderately methylated (10-50%; medium shade), and lowly methylated CpGs (< 10%; lightest shade) detected by each method **A)** for *M. capitata* and **B)** *P. acuta*, based on the number of CpGs captured by each method separately. Principal Coordinate Analyses associated with PERMANOVA and beta-dispersion tests related to Table ST6 that show differences in proportion of CpGs that are highly (≥ 50%), moderately (10-50%), or lowly (≤ 10%) methylated in **C)** *M. capitata* and **D)** *P. acuta*. WGBS is represented by green circles, RRBS by purple triangles, and MBDBS by orange diamonds. Percent variation explained by each PCoA axis is included in the axis label. Ellipses depict 95% confidence intervals for each sequencing method. All eigenvectors are significant at the α = 0.05 level.

**Figure SF4.jpg**

Methylation profile comparison of shared loci

PCA of CpG methylation for loci covered at 5x read depth in all samples for **A)** *M. capitata* and **B)** *P. acuta*.

**Figure SF5.jpg**

Genome coverage at various sequencing depths.

Estimated fraction of CpG sites in the genome covered by at least 5 reads at different sequencing depths (number of M read pairs) for **A)** *M. capitata* and **B)** *P. acuta* samples for each bisulfite sequencing method. ‘Observed’ (opaque line and dots) denotes the fraction of genome-wide CpG loci covered by at least 5 reads determined from pooled data that was subsampled at 50M, 100M, 150M, and 200M reads. ‘Estimated’ (translucent line and dots) denotes the fraction of genome-wide CpG loci covered by at least 5 reads estimated by michaelis-menten modelling of the ‘observed’ data with standard error shown by shaded areas. All samples within a bisulfite sequencing method were pooled for the downsampling analyses.

**Figure SF6.jpg**

Gene ontology enrichment of Biological Process terms from gene sets without CpG data from the three library prep methods in A) *M. capitata* and B) *P. acuta*. Colors indicate the overrepresented p values from GOseq analysis results with a p value cutoff of 0.01. See Table ST6,7,8,9 for full results with p values <0.05.

**Figure SF7.jpg**

Gene ontology enrichment of Molecular Function terms from gene sets without CpG data from the three library prep methods in A) *M. capitata* and B) *P. acuta*. Colors indicate the overrepresented p values from GOseq analysis results with a p value cutoff of 0.01. See Table ST6,7,8,9 for full results with p values <0.05.

**Figure SF8.jpg**

Coverage of orthologs across methods

Mean proportion (n=3 samples per method) of CpGs per gene that have at least 5x coverage in all of the one-to-one orthologous genes, as identified by OrthoFinder (Putnam et al., 2020) for **A)** *M. capitata* and **B)** *P. acuta*.

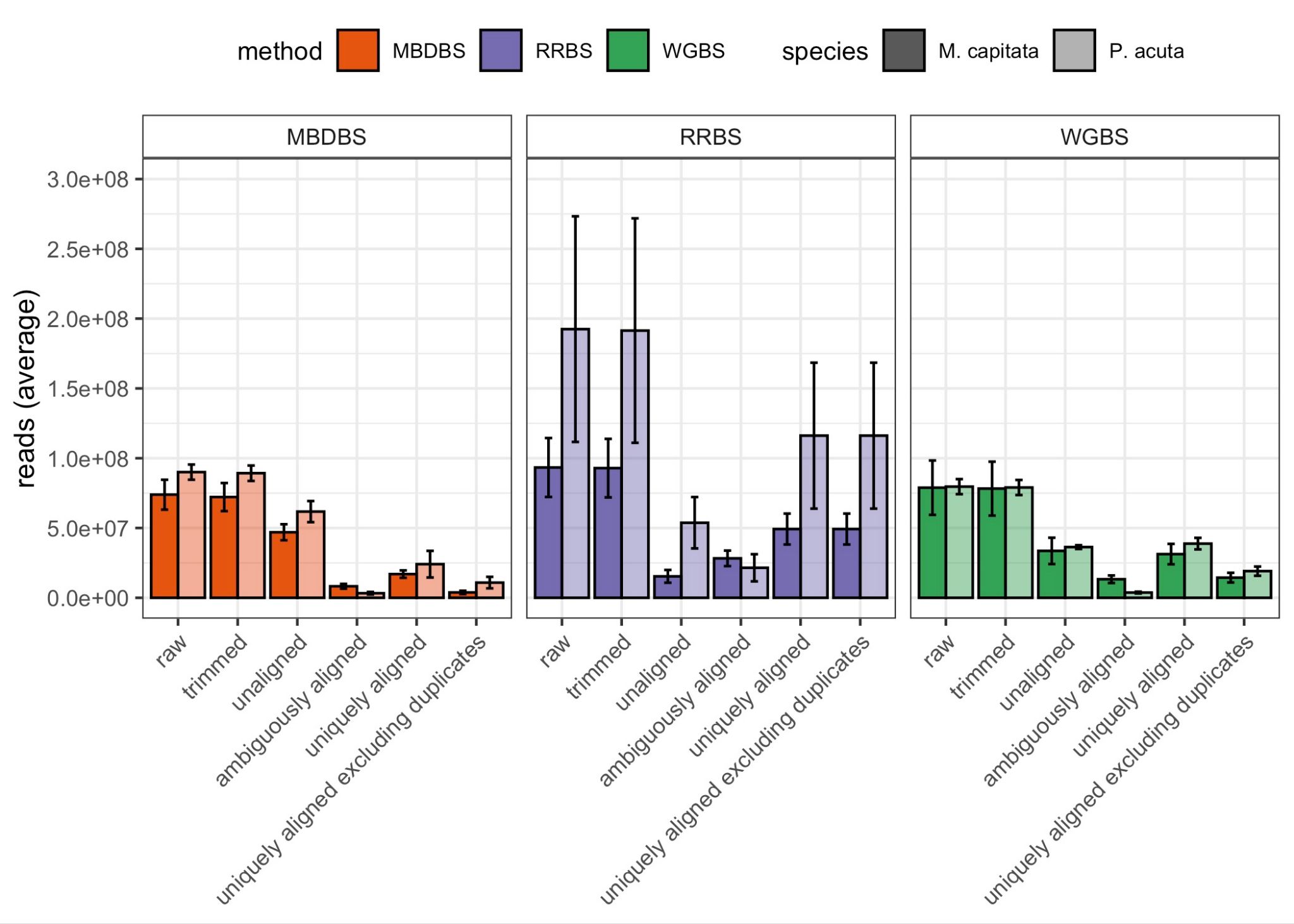

**Figure SF1.** Summary of sequencing depth and alignments for all libraries. Bars show average number of reads for each method and species and error bars show standard deviation.

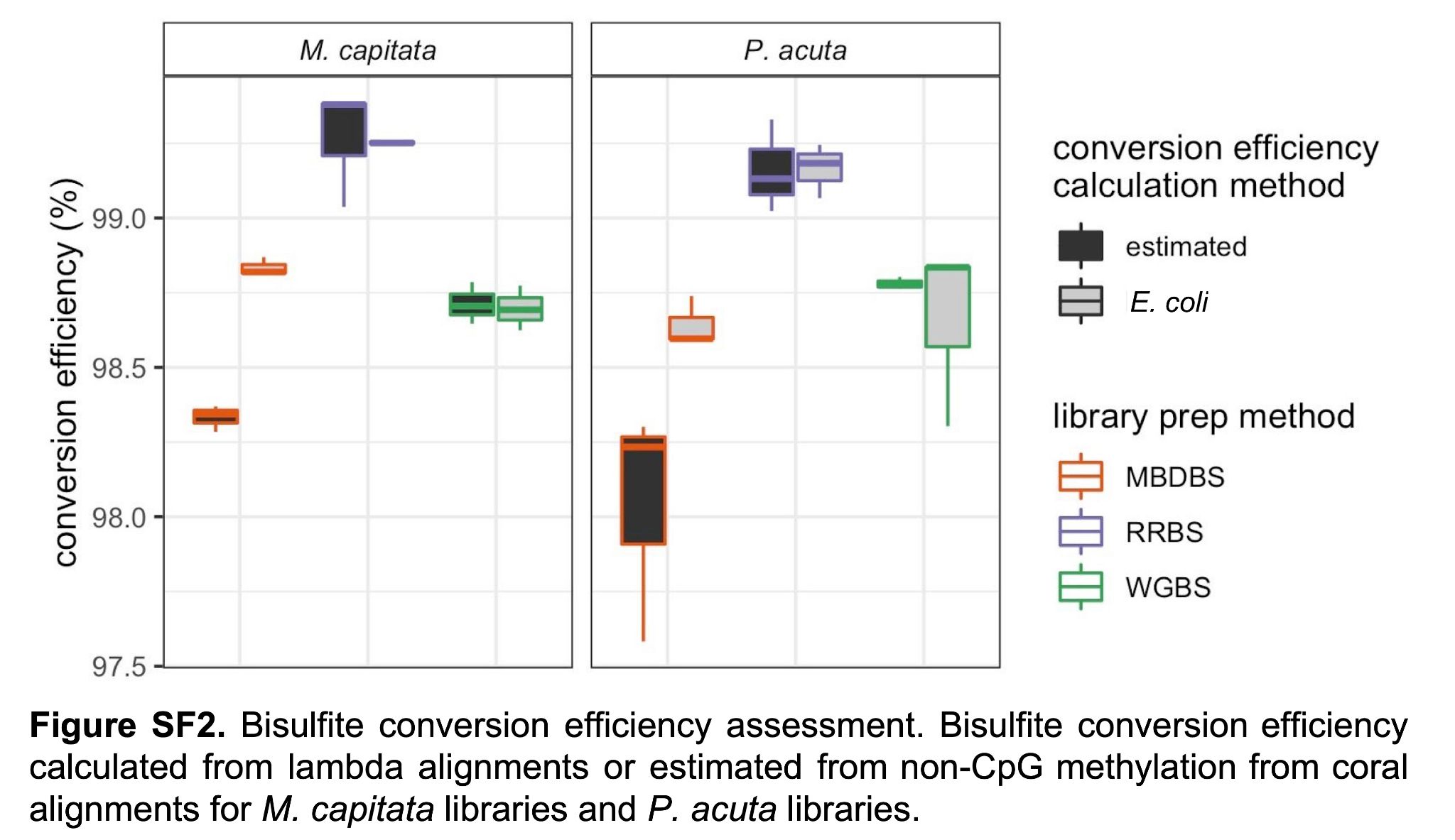

**Figure SF2.** Bisulfite conversion efficiency assessment. Bisulfite conversion efficiency calculated reads aligned to the *E. coli* unmethylated DNA spike, or estimated from non-CpG methylation from coral alignments for *M. capitata* libraries and *P. acuta* libraries.

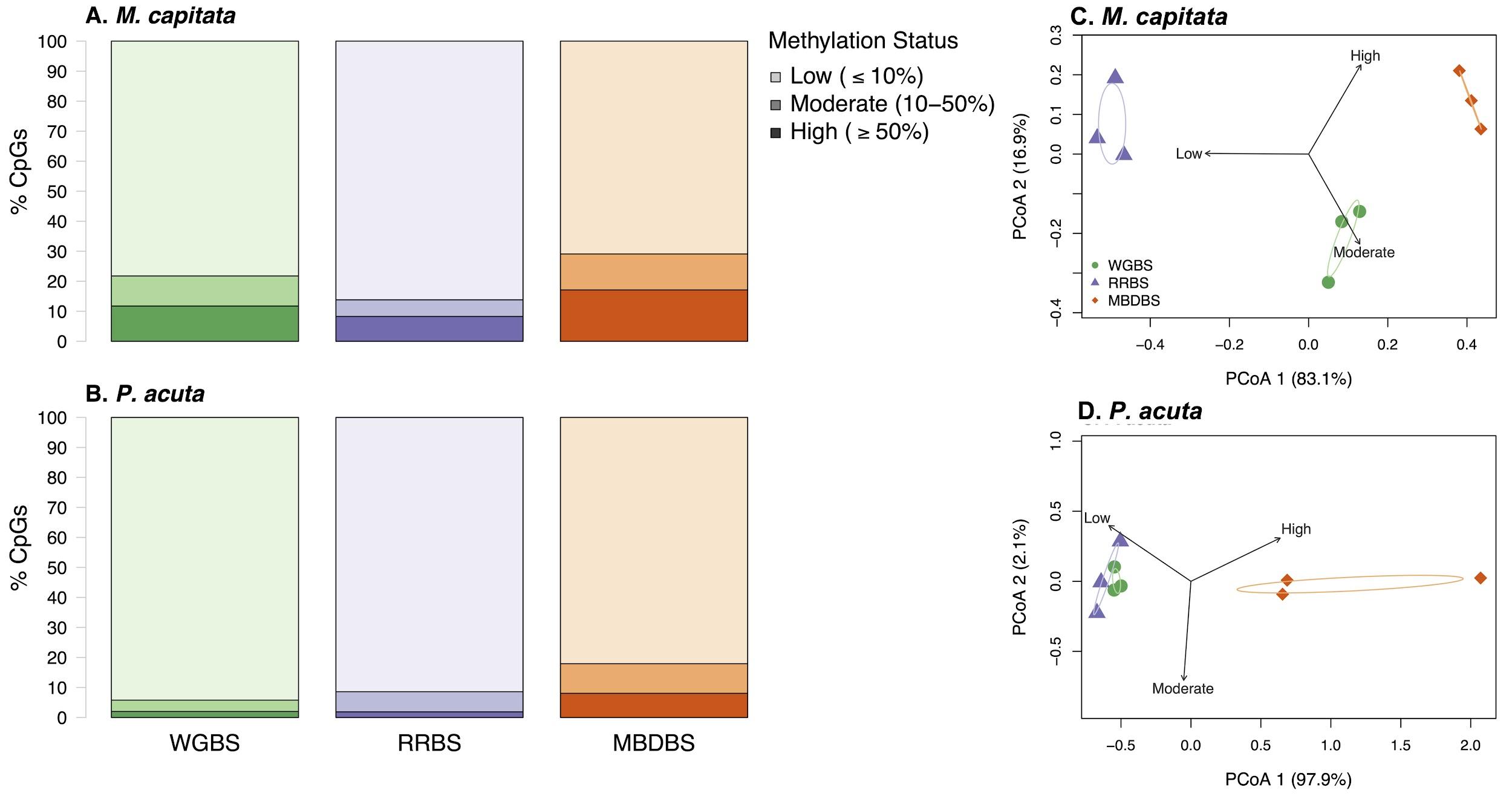

**Figure SF3**. Percent of highly methylated (≥ 50%; darkest shade), moderately methylated (10-50%; medium shade), and lowly methylated CpGs (< 10%; lightest shade) detected by each method **A**) for *M. capitata* and **B**) *P. acuta*, based on the number of CpGs captured by each method separately. Principal Coordinate Analyses associated with PERMANOVA and beta-dispersion tests related to **Table ST6** that show differences in proportion of CpGs that are highly (≥ 50%), moderately (10-50%), or lowly (≤ 10%) methylated in **C**) *M. capitata* and **D**) *P. acuta*. WGBS is represented by green circles, RRBS by purple triangles, and MBDBS by orange diamonds. Percent variation explained by each PCoA axis is included in the axis label. Ellipses depict 95% confidence intervals for each sequencing method. All eigenvectors are significant at the α = 0.05 level.

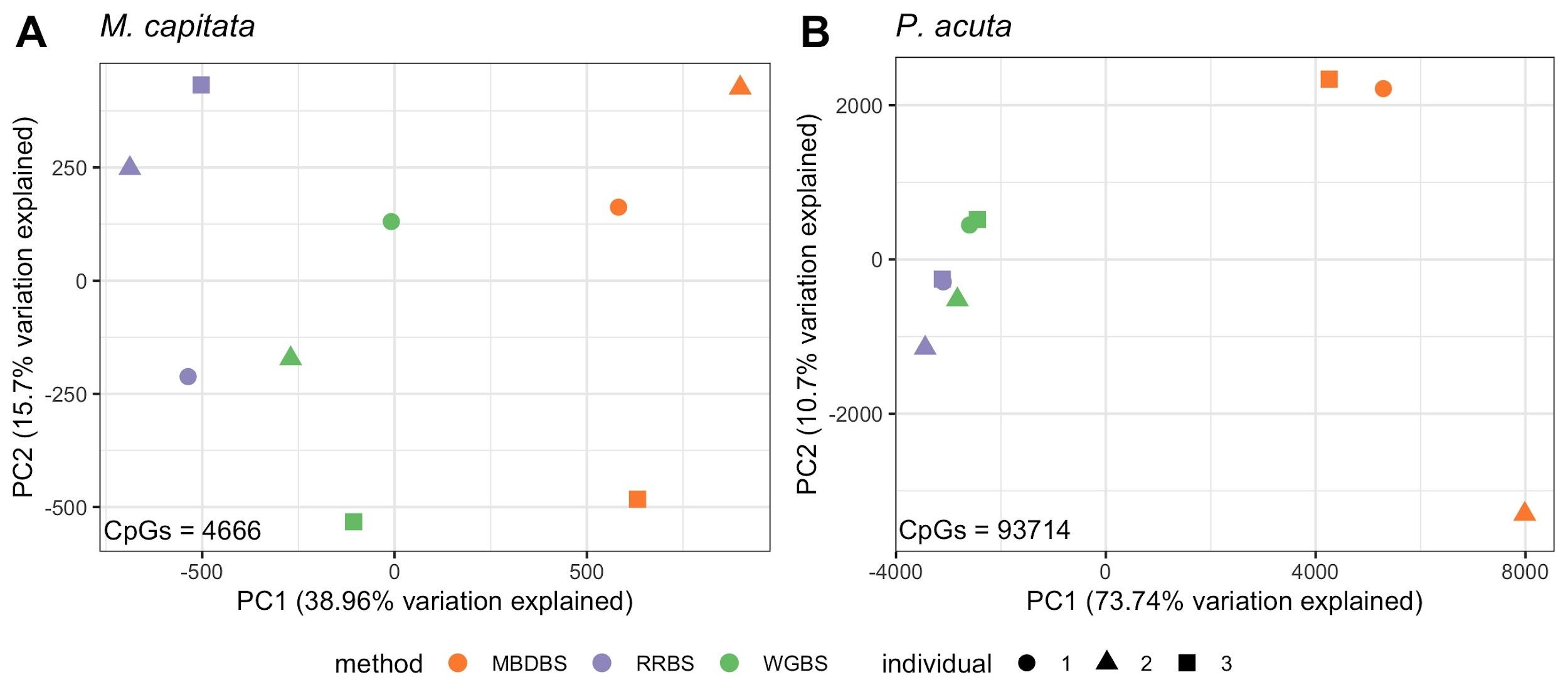

**Figure SF4.** PCA of CpG methylation for loci covered at 5x read depth in all samples for (**A**) *M. capitata* and (**B**) *P. acuta*.

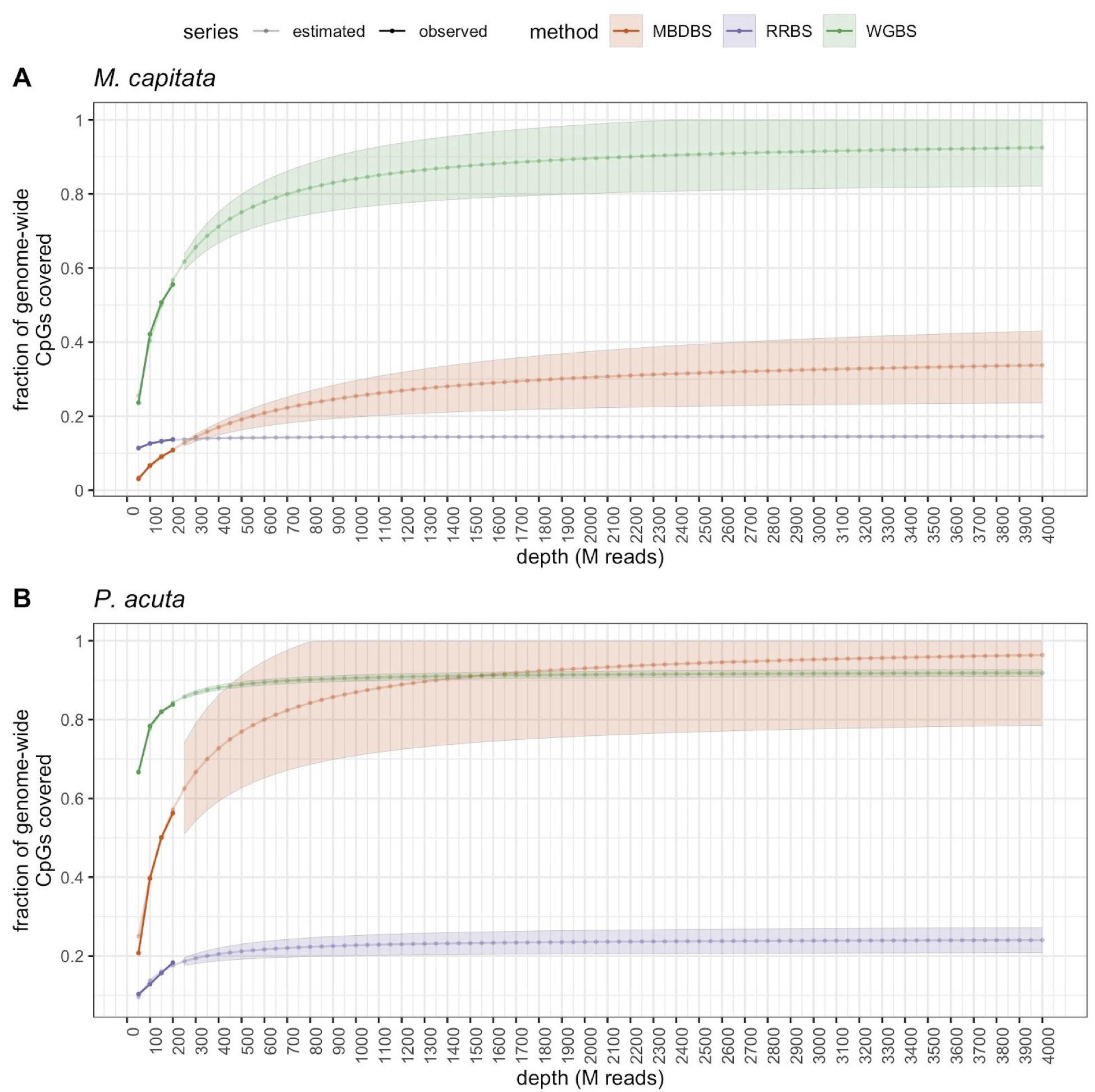
**Figure SF5.** Estimated fraction of CpG sites in the genome covered by at least 5 reads at different sequencing depths (number of M read pairs) for (**A**) *M. capitata* and (**B**) *P. acuta* samples for each bisulfite sequencing method. ‘Observed’ (opaque line and dots) denotes the fraction of genome-wide CpG loci covered by at least 5 reads determined from pooled data that was subsampled at 50M, 100M, 150M, and 200M reads. ‘Estimated’ (translucent line and dots) denotes the fraction of genome-wide CpG loci covered by at least 5 reads estimated by michaelis-menten modelling of the ‘observed’ data with standard error shown by shaded areas. All samples within a bisulfite sequencing method were pooled for the downsampling analyses.

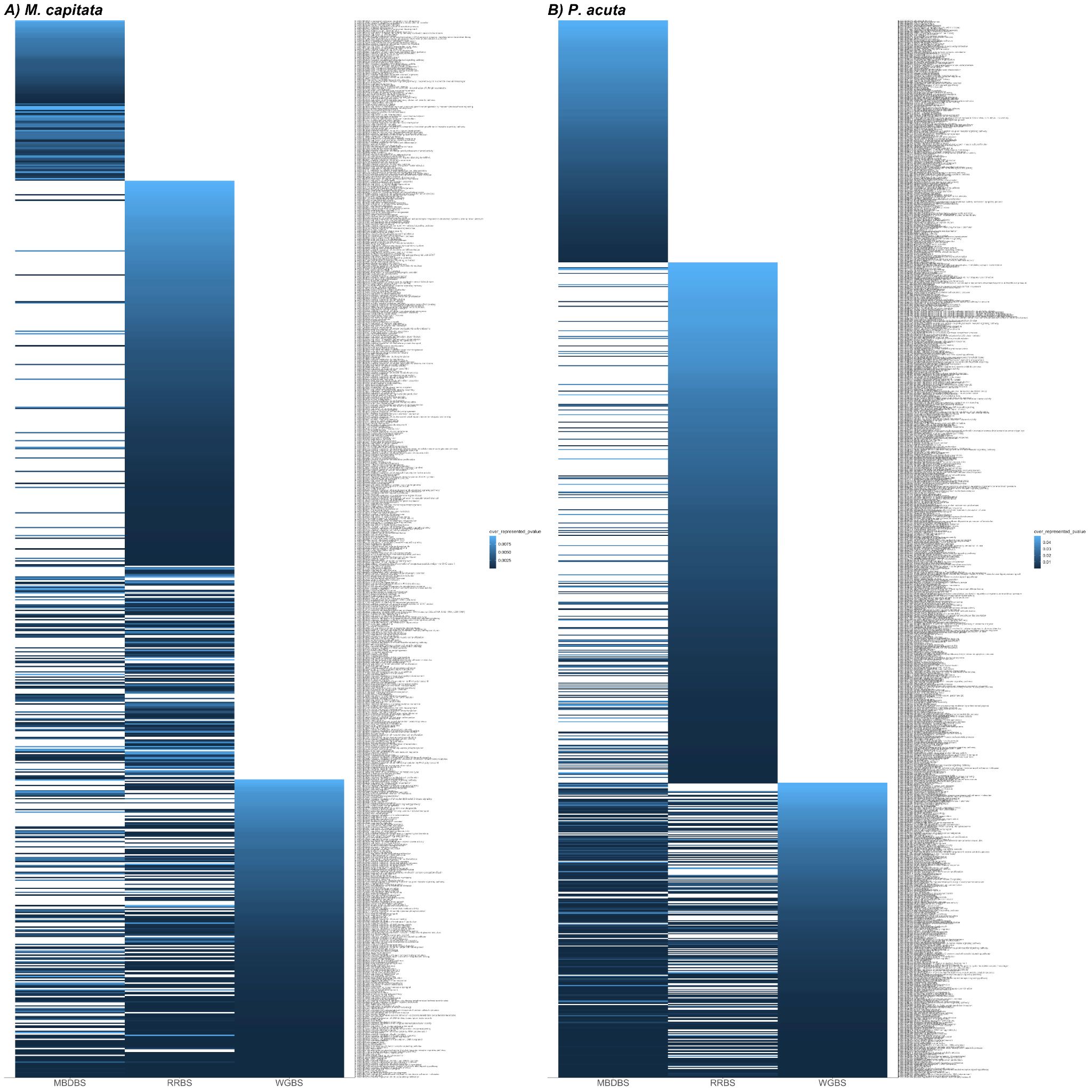

**Figure SF6.**

Gene ontology enrichment of Biological Process terms from gene sets with CpG data from the three library prep in A) *M. capitata* and B) *P. acuta*. Colors indicate the overrepresented p values from GOseq analysis results with a p value cutoff of 0.01. See Table ST6,7,8,9 for full results with p values <0.05.

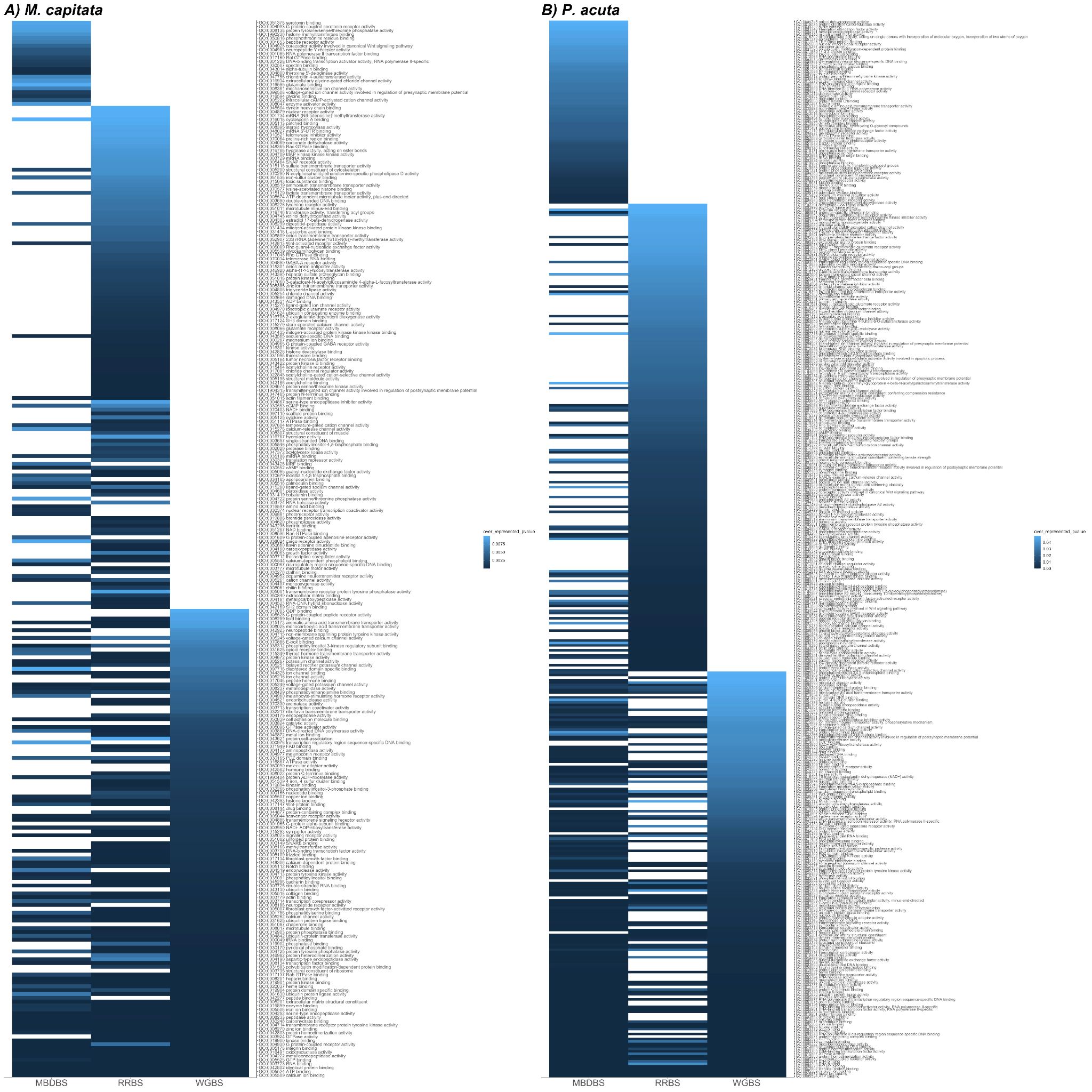

**Figure SF7.**

Gene ontology enrichment of Molecular Function terms from gene sets with CpG data from the three library prep methods in A) *M. capitata* and B) *P. acuta*. Colors indicate the overrepresented p values from GOseq analysis results with a p value cutoff of 0.01. See Table ST6,7,8,9 for full results with p values <0.05.

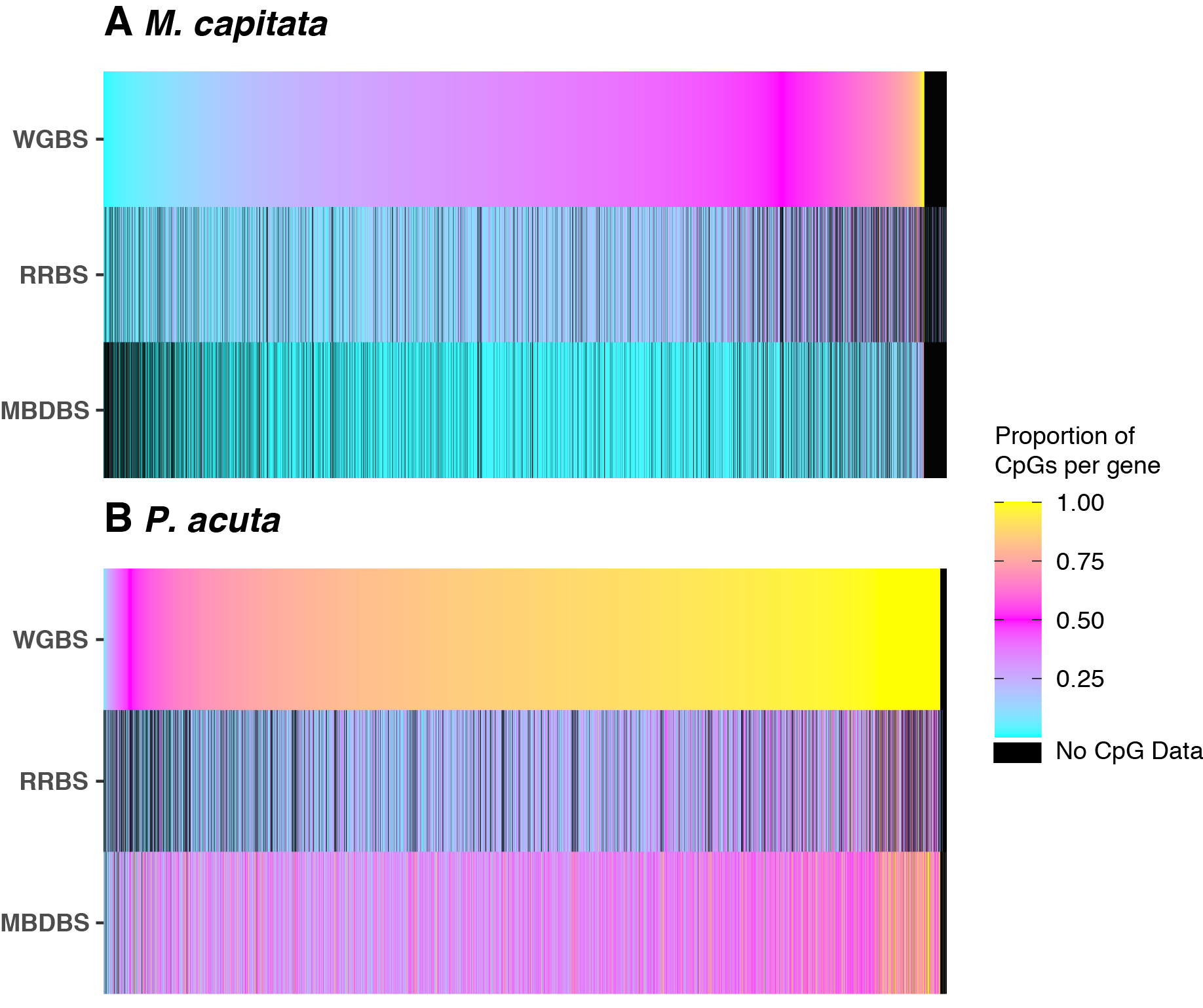

**Figure SF8.** Mean proportion (n=3 samples per method) of CpGs per gene that have at least 5x coverage in all of the one-to-one orthologous genes, as identified by OrthoFinder data available in [(Putnam, Roberts, and Venkataraman 2020)] for **A**) *M. capitata* and **B**) *P. acuta*.

Putnam, Hollie M., Steven B. Roberts, and Yaamini Venkataraman. 2020. “Coral Methylation Methods Comparison.” Open Science Framework. https://doi.org/[10.17605/OSF.IO/X5WAZ](http://dx.doi.org/10.17605/OSF.IO/X5WAZ).
